# The HDAC inhibitor trichostatin A impairs pancreatic β-cell function through an epigenome-wide reprogramming

**DOI:** 10.1101/2022.12.14.519294

**Authors:** Frédérik Oger, Maeva Moreno, Mehdi Derhourhi, Bryan Thiroux, Lionel Berberian, Cyril Bourouh, Emmanuelle Durand, Souhila Amanzougarene, Alaa Badreddine, Etienne Blanc, Olivier Molendi-Coste, Laurent Pineau, Gianni Pasquetti, Laure Rolland, Charlène Carney, Florine Bornaque, Emilie Courty, Céline Gheeraert, Jérôme Eeckhoute, David Dombrowicz, Julie Kerr-Conte, François Pattou, Bart Staels, Philippe Froguel, Amélie Bonnefond, Jean-Sébastien Annicotte

## Abstract

**Objective:** The pancreatic islets of Langerhans contain distinct cell subtypes including insulin-producing β cells. Although their cell-specific gene expression pattern defines their identity, the underlying molecular network driving this transcriptional specificity is not fully understood. Among the numerous transcriptional regulators, histone deacetylases (HDAC) enzymes are potent chromatin modifiers which directly regulate gene expression through deacetylation of lysine residues within specific histone proteins. The precise molecular mechanisms underlying HDAC effects on cellular plasticity and β-cell identity are currently unknown.

**Methods:** The pharmacological inhibition of HDAC activity by trichostatin A (TSA) was studied in the mouse Min6 and human EndocBH1 cell lines, as well as primary mouse sorted β cells and human pancreatic islets. The molecular and functional effects of treating these complementary β-cell models with TSA was explored at the epigenomic and transcriptomic level through next-generation sequencing of chromatin immunoprecipitation (ChIP) assays (ChIP-seq) and RNA sequencing (RNA-seq) experiments, respectively.

**Results:** We showed that TSA alters insulin secretion associated with β-cell specific transcriptome programming in both mouse and human β-cell lines, as well as on human pancreatic islets. We also demonstrated that this alternative β-cell transcriptional program in response to HDAC inhibition is related to an epigenome-wide remodeling at both promoters and enhancers.

**Conclusions:** Taken together, our data indicate that full HDAC activity is required to safeguard the epigenome, to protect against loss of β-cell identity with unsuitable expression of genes associated with alternative cell fates.

## 1. Introduction

The endocrine pancreas is composed of distinct cell subtypes including α (producing and secreting glucagon), β (producing and secreting insulin), δ (producing and secreting somatostatin), ε (producing and secreting ghrelin) and PP (producing and secreting pancreatic polypeptide) cells that play a crucial role in the regulation of glucose homeostasis [1]. A specific gene expression pattern defines each endocrine cell identity but the underlying molecular network that controls this transcriptional specificity remains elusive. The roles of some tissue-specific transcription factors, such as PDX1 or MAFA, in the maintenance of expression of genes controlling β-cell phenotype is well-known [2], but the contribution of chromatin modifiers in the maintenance of β-cell identity is less documented. Among these putative epigenomic regulators, there are histone acetyl transferases (HAT) and histone deacetylases (HDAC) that directly regulate gene expression through acetylation/deacetylation of lysine residues within specific histone and non-histone proteins [3].

HDACs are zinc metallo-enzymes divided into three main classes on the basis of their protein sequence homologies with yeast deacetylase enzymes [4]. Briefly, class I HDACs, composed of HDAC1, HDAC2, HDAC3 and HDAC8, are closely related to yeast Rpd3 (reduced potassium dependency 3) transcriptional regulator. Class II HDACs, including HDAC4, HDAC5, HDAC7, HDAC9 (class IIa) and HDAC6, HDAC10 (class IIb), share domains with similarity to yeast HdaI (histone deacetylase I), whereas HDAC11 belongs to the class IV. All these HDACs have a conserved catalytic domain and are therefore considered as ancestral enzymes that play a crucial role in the regulation of gene expression. While these enzymes have been considered as transcriptional inhibitors due to the resulting compaction of chromatin structure upon histone deacetylation [5], studies have also demonstrated that HDAC enzymes can actively contribute to cell-specific gene expression [6], suggesting that HDACs could play a dual active and inhibiting role in the regulation of gene expression to maintain cell identity.

The pharmacological inhibition of HDAC has gained a strong interest following the demonstration that HDAC inhibitors (HDACi) harbor anticancer properties [7]. HDACi suberoylanilide hydroxamic acid (SAHA, vorinostat) has been approved by the Food and Drug Administration for cancer therapy. HDACi may have potential as treatments for type 2 diabetes (T2D). Indeed, HDAC inhibition prevents cytokine-induced toxicity in β cells [8-10], and improves β-cell proliferation [11]. *Hdac3* β-cell specific knock down and pharmacological HDAC inhibition improve glucose tolerance [12; 13] and insulin sensitivity [14] in mice.

However, the treatment of the rodent β-cell line β-TC3 with HDACi also induces a loss of cell identity through a decrease of β-cell markers, correlated to an increase of α-cell marker expression within β cells [15], but the impact of HDACi on chromatin remodeling and the subsequent modulation of gene expression within β cells has not yet been interrogated.

Here, we explored the molecular and functional effects of treating β-cell models with the HDACi Trichostatin A (TSA) at the epigenomic and transcriptomic level. We show that TSA treatment leads to an epigenome-wide redistribution of histone marks that are enriched in active promoters and enhancers.

## 2. Materials and methods

### 2.1. Cell culture and chemicals

Chemicals, unless stated otherwise, were purchased from Sigma-Aldrich. Min6 cells (Addexbio) were maintained in DMEM high Glucose glutaMAX medium (Gibco, 31966-021) supplemented with 15 % SVF, 0.1 % β-mercaptoethanol and antibiotics (penicillin/streptomycin). Trichostatin A (Sigma, T1952-200UL) was used at different time and concentrations, as indicated. EndoC-βH1 cells were purchased from Human Cell Design™ and cultured according supplier recommendations.

Human pancreatic tissue was harvested from human, non-diabetic, adult donors. Isolation and pancreatic islet culture were performed as previously described [16].

### 2.2. Dot blot

Min6 cells were cultured in 24-well plates (2.10^5^ cells/well) and were resuspended in 100 μL of RIPA buffer (Thermo Scientific, 89901), then incubated at 4 °C during 30 min under agitation. After centrifugation (15000 g, 20 min, 4 °C), supernatant was harvested and protein concentration was measured using BCA assay (Pierce, 23227). Samples were denatured (95 °C, 5 min) and 1 μg of total protein were dotted on nitrocellulose membrane using a dot blotter (Cleaver Scientific Ltd, CSL-D96). After drying (15 min, RT), membranes were saturated in saturation buffer (5 % free fatty acids milk in TBS 1X, 2h, room temperature), then incubated with primary antibody diluted in saturation buffer (16 h, 4 °C). After incubation with HRP-coupled secondary antibody diluted in saturation buffer (1 h, room temperature), membranes were revelated using ECL kit (Pierce, 34076) on Chemidoc XRS+ imager (Biorad). Images were processed and analyzed using ImageJ software. The list of antibodies used in this study is available in the Supplementary Table S1.

### 2.3. Animal procedures

Mice were maintained according to European Union guidelines for the use of laboratory animals. *In vivo* experiments were performed in compliance with the French ethical guidelines for studies on experimental animals (animal house agreement no. A59-35015, Authorization for Animal Experimentation, project approval by our local ethical committee no. APAFIS#2915-201511300923025v4). C57bl6J (Charles River Laboratories) and Rip-Cre::Lox-STOP-Lox-Tomato (obtained after breeding of Rip-Cre mice [17] with LSL-Td-Tomato (Jax, #stock number 007905)) mice were maintained under 12 hours light/dark cycle and were fed *ad libitum*.

### 2.4. Mouse pancreatic islet isolation and cell sorting

Mouse pancreatic islets were prepared as described previously [18]. Briefly, pancreata were digested using type V collagenase (Sigma, ref C9263-1G) for 10 min at 37 °C. After digestion, pancreatic islets were separated in a density gradient medium (Histopaque) and purified by handpicking under a macroscope. Pancreatic islets were cultured for 24 hours before further processing. For cell sorting experiments, mouse pancreatic islets isolated from RipCre-tdTomato mice were trypsinized for 7 minutes with 1 mM of trypsine (Gibco). Dissociated pancreatic cells were directly run into an Influx sorter (Becton Dickinson®) equipped with a 86 μm nozzle and tuned at a pressure of 24.7 psi and a frequency of 48.25 kHz. Sample fluid pressure was adjusted to reach and event rate of 10 000 events/second. β cells and other pancreatic cells were selected as Tomato + (Tom+, β cells) and Tomato – (Tom-, non-β cells) and sorted in purity mode (phase mask 16/16) directly in RNeasy kit Lysis buffer for further processing.

### 2.5. Glucose stimulated insulin secretion (GSIS)

Min6 cells cultured in 96-well plates (2.10^4^ cells/well) were glucose-starved in 200 μL of starvation buffer (Krebs Ringer buffer (KRB) supplemented with BSA 0.5 %, 1 h, 37 °C, 5 % CO_2_). After incubation, the starvation buffer was discarded and cells were incubated in 200 μL of 2.8 mM glucose-concentrated starvation buffer (1 h, 37 °C, 5 % CO_2_). After 2.8 mM glucose samples recovering, cells were incubated in 200 μL of 20 mM glucose-concentrated starvation buffer (1 h, 37 °C, 5 % CO_2_). After 20 mM glucose samples recovering, the intracellular insulin content was recovered in 100 μL of lysis buffer (Ethanol 75 %, HCl 1.5 %). Insulin concentration was measured using mouse or human Insulin ELISA kits according the manufacturer’s instructions (Mercodia).

### 2.6. RNA extraction and RT-qPCR

Total RNA was extracted from Min6 cells using RNeasy Plus mini kit and from pancreatic islets using RNeasy Plus micro kit (Qiagen, ref 74034) following the manufacturer’s instruction. Total RNA sample concentrations were determined using a Nanodrop spectrophotometer (Implen). 100 ng of RNA was used for reverse transcription (RT) using Transcriptor universal cDNA master mix (Roche) for Min6 cells or superscript III enzyme (Invitrogen) for mouse pancreatic islets according to the manufacturer’s instructions. Gene expression was measured through quantitative real-time PCR (qPCR) using FastStart SYBR Green master mix (Roche) according to the manufacturer’s recommendations in a LightCycler 480 II device (Roche). Mouse RT-qPCR results were normalized to endogenous cyclophilin reference mRNA levels. The results are expressed as the relative mRNA level of a specific gene expression using the formula 2^−ΔΔCt^. The oligonucleotides sequences used for various experiments are listed in Supplementary Table S2.

### 2.7. RNA sequencing (RNA-seq)

RNA quality was verified using RNA 6000 nanochips (Agilent, #5067-1511) on the Agilent 2100 bioanalyzer (Agilent, #G2939A). 500 ng of purified RNA with RNA integrity number (RIN) ≥8 was subsequently used for library preparation with the TruSeq Stranded mRNA library Prep (Illumina, #20020594) and sequenced on the Illumina NextSeq500 system using a paired-end 2×75 bp protocol. Raw sequencing data are available upon request.

### 2.8. Chromatin immunoprecipitation sequencing (ChIP-seq)

20.10^6^ Min6 cells were treated with formaldehyde at a final concentration of 1% to cross link DNA and protein complexes during 10 min. The reaction was stopped by the addition of Glycine 0.125 M during 5 min. Cells were lysed and DNA-protein complexes were sheared using the Bioruptor Pico (Diagenode, #B01060010) for 8 minutes. The sheared chromatin was followed by immunoprecipitation with either the non-specific antibody IgG (Santa Cruz, #sc2025), H3K4me1 (Active motif, #39297), H3K4me3 (Active motif, #61379), H3K27me3 (Active motif, #61017), H3K27ac (Active motif, #39685) or H3K9ac (Abcam, #ab47915). 1 ng of eluted and purified DNA was used to prepare libraries with the Nextflex rapid DNA seq kit 2.0 (Perkin Elmer, #NOVA-5188-01) on the Illumina NextSeq500 system using a single read 100 bp protocol. Raw sequencing data are available upon request.

### 2.9. Bioinformatic analysis

#### 2.9.1. RNA-seq

The demultiplexing of sequence data (from BCL files generated by Illumina sequencing systems to standard FASTQ file formats) was performed using bcl2fastq Conversion Software (Illumina; version 2.19.1). Trimming of residuals adapters and low-quality reads was performed using Cutadapt software (version 1.7.1). Subsequently, sequence reads from FASTQ files were mapped to the mouse genome (mm10) using STAR Aligner (version 2.5.2b). On average, 38M reads were generated per sample, and 93.5 % +/- 0.9% of them were accurately mapped. The counting of the different genes and isoforms was performed using RSEM (version 1.3). Finally, differential expression was performed using DESeq2 package.

#### 2.9.2. ChIP-seq

The demultiplexing of sequence data (from BCL files generated by Illumina sequencing systems to standard FASTQ file formats) was performed using bcl2fastq Conversion Software (Illumina; version 2.20). Trimming of residuals adapters and low-quality reads was performed using TrimGalore (version 0.4.5). Subsequently, sequence reads from FASTQ files were mapped to the mouse genome (mm10) using Bowtie2 Aligner (version 2.3.5.1). Finally peak-calling was performed using MACS2 software (version 2.2.7.1).

#### 2.9.3. Pathway analysis

RNA-seq data and integrated ChIP-seq/RNA-seq data were uploaded to Metascape website [19] or Ingenuity Pathway Analysis software (Qiagen) as specified. For RNA-seq data, an adjusted p-value < 0.05, LogFC>1 and LogFC<-1 were set as thresholds as indicated and pathway analyses were performed using the core analysis function, including canonical pathways, upstream regulators, diseases, biological functions and molecular networks.

### 2.10. Statistical analysis

Data are presented as mean ± s.e.m. Statistical analyses were performed using a two-tailed unpaired Student’s t-test, one-way analysis of variance (ANOVA) followed by Dunnett’s post hoc test or two-way ANOVA with Tukey’s post hoc tests comparing all groups to each other, using GraphPad Prism 9.0 software. Differences were considered statistically significant at p < 0.05 (*p < 0.05, ** p < 0.01, *** p < 0.001 and **** p < 0.0001).

## 3. Results

### 3.1. TSA-mediated HDAC inhibition increases global histone acetylation associated to impaired insulin secretion and decreased expression of β-cell identity genes

We first characterized the role of HDAC inhibition in Min6 cells through a pharmacological approach using the pan-HDAC inhibitor TSA. A time-course TSA treatment (0.5 μM) of Min6 cells was performed from 15 min to 24 h and acetylation of lysine 27 (H3K27ac), lysine 9 (H3K9ac) and global acetylation (PanH3ac) levels of histone H3 were monitored by dot blot to validate global HDAC inhibition (Figure 1A). TSA induced a time-dependent increase of H3K27ac (Figures 1A and B), H3K9ac (Figures 1A and C) and PanH3ac (Figures 1A and D), indicating that Min6 cells were sensitive to HDAC inhibitor treatment. Given that the sub-maximal histone H3K27 and H3K9 acetylation signal was observed 16 h after TSA treatment (Figures 1A, B and C), subsequent experiments were performed at this time point. To ensure that TSA did not induced cytotoxicity, the viability of Min6 cells upon 16 h of TSA treatment was assessed by FACS through annexin V and propidium iodide labeling to identify both apoptosis events and cell viability, respectively. Neither annexin V (Supplementary Figure S1A) nor propidium iodide (Supplementary Figure S1B) positive cells were significantly overrepresented upon TSA treatment compared to vehicle (DMSO 0.1%) treatment, indicating that TSA did not affect Min6 cell viability. Then, to determine the effects of HDAC inhibition on specific acetylation and methylation sites of histone H3 involved in transcriptional regulation, the acetylation level of lysine 9 and 27 of histone 3 (H3K9ac and H3K27ac, respectively) and the trimethylation level of lysine 27 (H3K27me3) were specifically monitored by dot blot in Min6 cells after 16h of TSA treatment. Compared to vehicle-treated cells, TSA induced a significant increase of both H3K9ac (Figures 1E and F) and H3K27ac (Figure 1E and G) indicating that the effects of HDAC inhibition in Min6 cells could be partly related to the increased acetylation of these histone marks. Trimethylation level of lysine 27 of H3 (H3K27me3) was significantly decreased upon TSA treatment (Figure 1E and H). Then, to evaluate the impact of TSA on functional properties of Min6 cells, insulin secretion was measured through glucose-stimulated insulin secretion (GSIS) assays. TSA treatment significantly increased insulin secretion in low glucose conditions (2.8 mM) and significantly decreased insulin secretion in high glucose conditions (20 mM, Figure 1I). These functional effects were associated with a decreased expression of genes involved in β-cell functions, such as *Pdx1* or *Mafa* (Figure 1J). These results suggest that HDACs maintain functional properties of β cells through mechanisms that imply the modulation of histone acetylation and gene expression.

**Figure 1.**
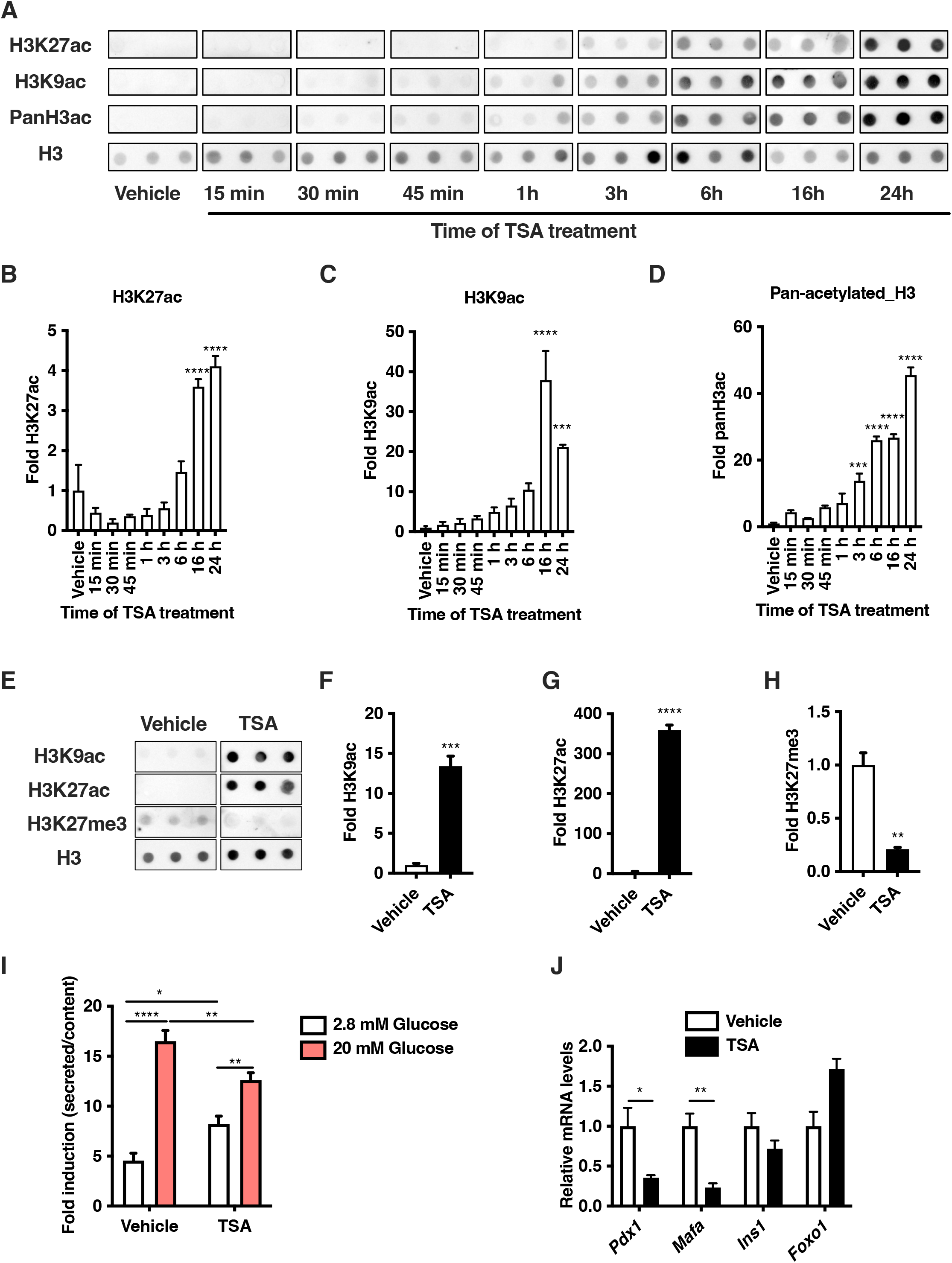
The increase of histone H3 acetylation level upon TSA treatment is correlated to alteration of functional properties of Min6 cells. **(A)** Dot blot kinetic analysis of H3K9ac, H3K27ac and pan-acetylated histone H3 (Pan H3ac) levels upon TSA treatment (0.5 μM) in Min6 cells at the indicated time. Vehicle (DMSO 0.1%) was used as negative control. **(B-D)** Densitometry analysis of A was performed by comparing the mean of signal at each time point to mean signal in vehicle (n=3). **(E)** Dot blot analysis of H3K9ac, H3K27ac and H3K27me3 levels upon TSA treatment (0.5 μM, 16h) in Min6 cells. Vehicle (DMSO 0.1%) was used as negative control. **(F-H)** Densitometry analysis of E is expressed as fold of H3K9ac (F), H3K27ac (G) and H3K27me3 (H) signals in TSA-treated cells compared to signals in vehicle-treated cells (n=3). **(I)** Glucose-stimulated insulin secretion of TSA-treated cells (0.5 μM, 16h, n=7). Vehicle (DMSO 0.1%) was used as a control. Results are presented as fold insulin secretion in response to 20 mM glucose compared to 2.8 mM glucose +/- SEM. **(J)** Quantitative RT-PCR analysis of key β-cell identity genes in vehicle and TSA-treated Min6 cells (n=3). Results in B, C, D, F, G, H, I, J are displayed as means +/- SEM. * p<0.05, ** p<0.01, *** p<0.001, **** p<0.0001.

### 3.2. Epigenome-wide histone mark and transcriptome profiling identifies distinct functional genomic regions in Min6 cells

With the aim to restrict our analysis to the effects of HDAC inhibition on functional genomic regions, we first performed profiling of H3K4me3, H3K4me1, H3K27ac and H3K27me3 through ChIP-seq experiments in untreated, control Min6 cells to delineate these specific chromatin features in basal conditions. As expected, H3K4me3 and H3K27ac were mostly enriched within gene promoter (74% and 58%, respectively, Supplementary Figures 2A and 2B) whereas H3K4me1 and H3K27me3 were mostly enriched both within gene body (46% and 40%, respectively) and intergenic regions (25% and 44%, respectively, Supplementary Figures 2C and 2D). Interestingly, this systematic histone mark profiling at the epigenome-wide level allowed us to precisely circumscribe 6 distinct functional genomic regions based on combinatorial histone mark enrichment within both promoter and distal intergenic regions. Indeed, at the promoter level, we defined active (*i.e*., both H3K4me3 and H3K27ac enriched; Figure 2A), bivalent (*i.e*., both H3K4me3 and H3K27me3 enriched; Figure 2B) and inactive (*i.e*., no histone marks enrichment; Figure 2C) promoters. Within distal intergenic regions, we were able to define active enhancers (*i.e*., both H3K27ac and H3K4me1 enriched; Figure 2D), poised enhancers (*i.e*., both H3K4me1 and H3K27me3 enriched; Figure 2E) and heterochromatin (*i.e*., only H3K27me3 enriched; Figure 2F).

**Figure 2.**
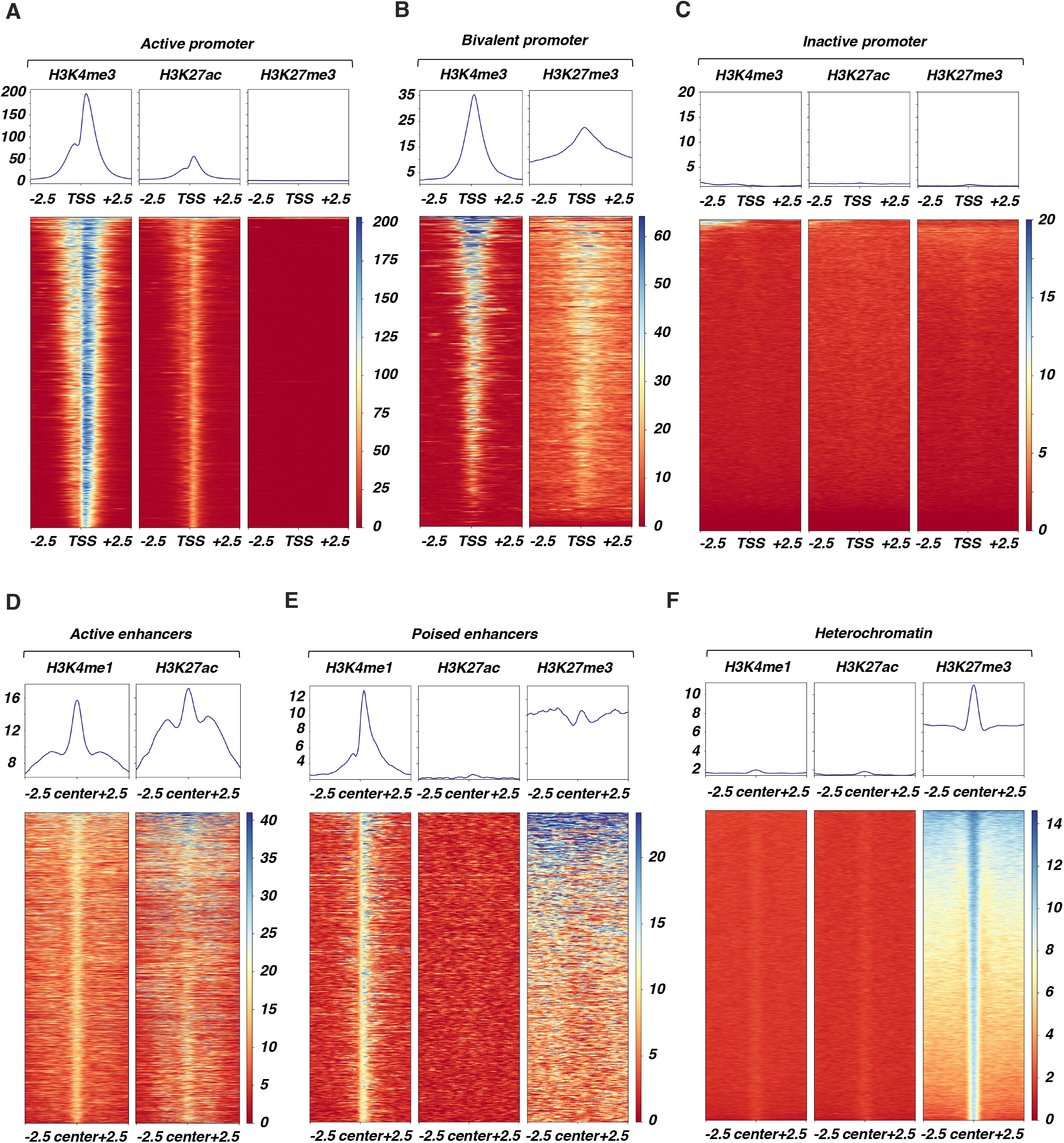
Histone marks profiling by ChIP-seq defines distinct functional genomic regions in Min6 cells. **(A)** Characterization of active promoters in Min6 cells. Heatmap displayed signal centered on transcription start site (TSS) +/- 2.5kb for H3K4me3, H3K27ac and H3K27me3 within each active promoter (n=8303 regions, 2549 genes). Mean signal centered on TSS +/- 2.5kb for H3K4me3, H3K27ac and H3K27me3 within active promoters is displayed. **(B)** Characterization of bivalent promoters in Min6 cells. Heatmap displayed signal centered on TSS +/- 2.5kb for H3K4me3 and H3K27me3 within each bivalent promoter (n=3685 regions, 1323 genes). Mean signal centered on TSS +/- 2.5kb for H3K4me3 and H3K27me3 within bivalent promoters is displayed. **(C)** Characterization of inactive promoters in Min6 cells. Heatmap displayed signal centered on TSS +/- 2.5kb for H3K4me3, H3K27ac and H3K27me3 within each inactive promoter (n=7632 genes). Mean signal centered on TSS +/- 2.5kb for H3K4me3, H3K27ac and H3K27me3 within inactive promoters is displayed. **(D)** Characterization of active enhancers in Min6 cells. Heatmap displayed signal centered on the center of the regions +/- 2.5kb for H3K4me1 and H3K27ac within each active enhancer (n=1597 regions). Mean signal centered on the center of the regions +/- 2.5kb for H3K4me1 and H3K27ac within active enhancers is displayed. **(E)** Characterization of poised enhancers in Min6 cells. Heatmap displayed signal centered on the center of the regions +/- 2.5kb for H3K4me1, H3K27ac and H3K27me3 within each poised enhancer (n=683 regions). Mean signal centered on centered on the center of the regions +/- 2.5kb for H3K4me1, H3K27ac and H3K27me3 within poised enhancers is displayed. **(F)** Characterization of heterochromatin in Min6 cells. Heatmap displayed signal centered on the center of the regions +/- 2.5kb for H3K4me1, H3K27ac and H3K27me3 within heterochromatin (n=21292 regions). Mean signal centered on centered on the center of the regions +/- 2.5kb for H3K4me1, H3K27ac and H3K27me3 within heterochromatin is displayed.

To go further in the characterization of these distal functional intergenic regions, their associated genes were defined using Genomic Regions Enrichment of Annotations Tool (GREAT) software [20].

Secondly, with the aim to link these chromatin features to the associated-gene expression level, transcriptomic analysis through RNA-seq was performed in untreated Min6 cells. The genes associated with active regions such as active promoter as well as active enhancers were significantly found to be the most expressed compared to genes associated with other chromatin features (*i.e*., bivalent and inactive promoter, poised enhancers and heterochromatin, Figure 3A). The expression level of genes associated with poised enhancers was not significantly different to the expression level of genes associated with heterochromatin (Figure 3A). Then, we analyzed the gene ontology of genes associated with these functional genomic regions. This analysis showed that biological processes related to genes associated with functionally active genomic regions were enriched either in pathways involved in general biological functions such as mitotic cell cycle process (-Log10(q)=22, active promoter, Figure 3B) or in β-cell specific functions such as the regulation of secretion ((-Log10(q)=34, active enhancers, Figure 3C). Conversely, genes associated with inactive bivalent, poised genomic regions were rather enriched in pathways unrelated to β-cell functions such as embryonic morphogenesis (-Log10(q)=35, bivalent promoters, Figure 3D; - Log10(q)=325, heterochromatin, Figure 3F), dorsal spinal cord development (-Log10(q)=19, poised enhancers, Figure 3E) or cell fate commitment (-Log10(q)=29, bivalent promoters, Figure 3D; -Log10(q)=325, heterochromatin, Figure 3F). Due to the large number of genes associated with inactive promoters in Min6 cells, no specific pathway was identified for these associated genes (data not shown). Altogether, these results indicate that, based on the combinatorial enrichment of a selection of histone marks involved in transcriptional regulation in Min6 cells, these chromatin features were functional and directly related to gene expression level and the regulation of biological process involved in β-cell function.

**Figure 3.**
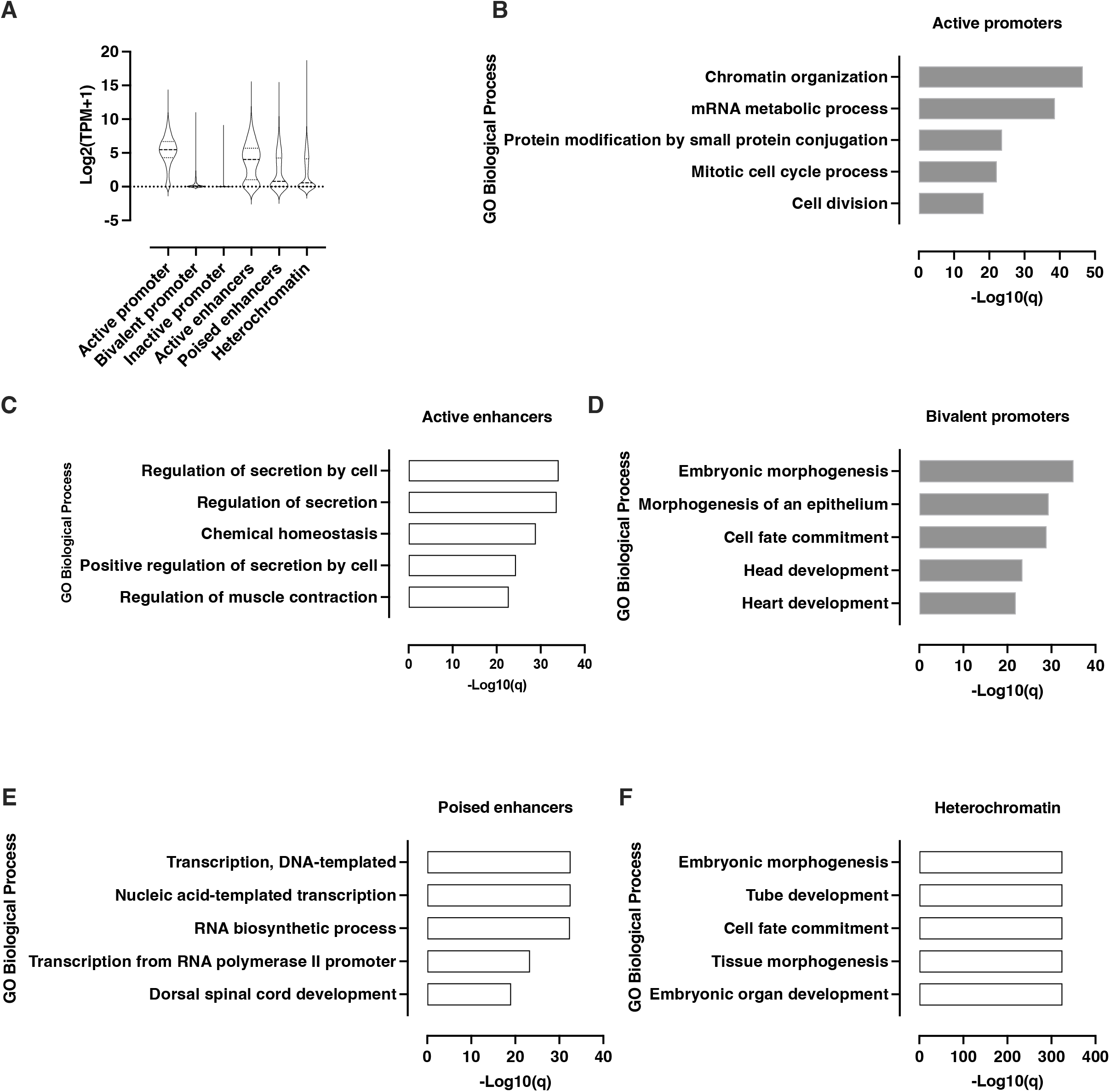
Functional genomic regions in Min6 cells are directly associated with gene expression level and distinct biological processes. **(A)** RNA-seq based expression level of genes associated with functional genomic regions in Min6 cells. Results are displayed as mean of Log2(TPM+1) for each functional genomic regions associated-genes. The violin plot displays median and quartile values for each group of genes. **(B)** Gene ontology (GO) analysis of active promoter associated-genes was performed using Metascape by filtering output only on GO Biological process. The 5 more enriched GO Biological processes are displayed as -Log10(q) (*i.e*., - Log10(Padj)). **(C)** GO was performed using Genomic Regions Enrichment of Annotations Tool (GREAT) following the two nearest genes association rule (cut-off<1000kb). The 5 more enriched GO biological processes are displayed as -Log10(q) (*i.e*., -Log10(Padj)). **(D)** GO analysis was performed using Metascape by filtering output only on GO Biological process. The 5 more enriched GO Biological processes are displayed as -Log10(q) (*i.e*., -Log10(Padj)). **(E)** GO analysis was performed using GREAT following the two nearest genes association rule (cut-off<1000kb). The 5 more enriched GO biological processes are displayed as -Log10(q) (*i.e*., -Log10(Padj)). **(F)** GO analysis was performed using GREAT following the two nearest genes association rule (cut-off<1000kb). The 5 more enriched GO biological processes are displayed as -Log10(q) (*i.e*., -Log10(Padj)).

### 3.3. The functional genomic regions identified in Min6 cells overlap with functional chromatin segments found in mouse islets

To go further in the characterization of these functional genomic regions in Min6 cells, we next assessed whether they overlapped with functional genomic regions recently identified in mouse islets [21]. Therefore, we first sought to determine the overlapping level between the genomic regions (peaks) independently enriched in a selection of histone marks in Min6 cells and the functional genomic segments resulting from functional chromatin segmentation (categorized from A to X) defined in mouse islets of Langerhans [21]. This analysis showed that most of the regions enriched in H3K4me3 in Min6 cells were mostly found in active (segments A to D) or bivalent (segment K to M) chromatin segments, suggesting that the active promoter regions of mouse β cells are conserved in Min6 cells (Figure 4A, H3K4me3). This result was corroborated by data obtained with the genomic regions enriched in H3K27ac in Min6 cells showing that these also mostly overlapped with the segments of active chromatin (segments A to D, Figure 4A, H3K27ac). Furthermore, the regions enriched with H3K4me1 in Min6 cells were also conserved since they predominantly overlapped with the active chromatin segments (segments from C to E) and to a lesser extent with the bivalent regions (segment K, Figure 4A, H3K4me1). Regarding the regions enriched in H3K27me3 in Min6 cells, these were mainly found in the segments of silent chromatin (segments N to P, Figure 4A, H3K27me3). Taken together, this analysis demonstrated that the regions enriched in histone marks involved in transcriptional regulation are, at least partially, shared between Min6 cells and mouse pancreatic islets.

**Figure 4.**
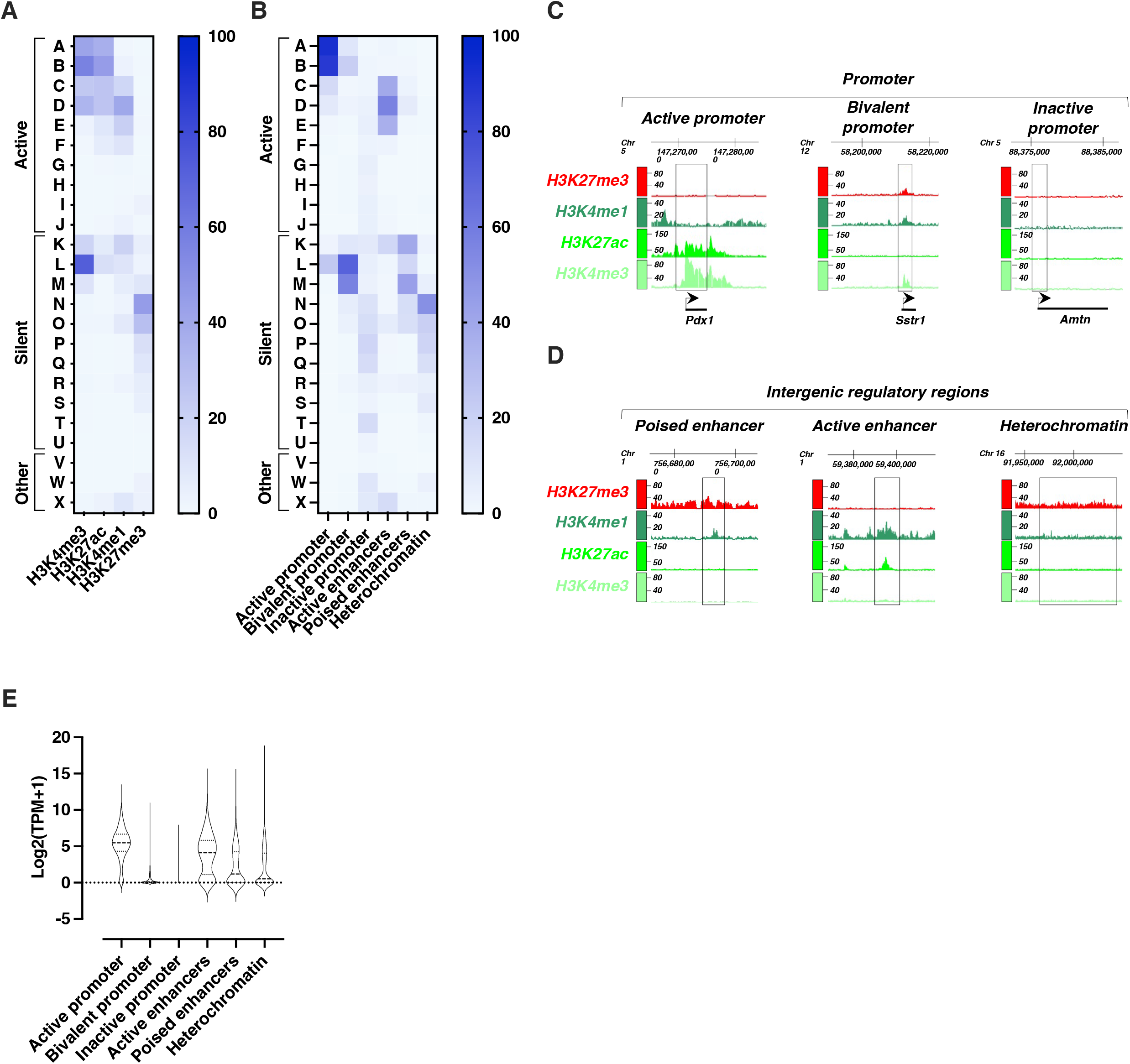
Functional genomic regions in Min6 cells are partly conserved with functional genomic segments in mouse islets of Langerhans. **(A)** Percentage overlap of H3K4me3, H3K27ac, H3K4me1 and H3K27me3 peaks in Min6 cells with the distinct genomic segments (A to J, active segments; K to U, silent segments; V to X, others segments) from mouse pancreatic islets chromatin segmentation [21]. Peaks for each histone mark were intersected with each mouse islets chromatin segment to define the overlap associated-percentage. Results are displayed as heatmap scaled on this percentage of overlap. **(B)** Percentage overlap of functional genomic regions in Min6 cells with the distinct genomic segments from mouse islets of Langerhans chromatin segmentation [21]. Each specific functional genomic region was intersected with each mouse pancreatic islets chromatin segment to define the overlap associated-percentage. Results are displayed as heatmap scaled on this percentage of overlap. **(C-D)** Examples of conserved functional genomic regions. Figures are adapted from the IGB genome browser screenshots and genomic coordinates are indicated. **(E)** RNA-seq based expression level of genes associated with conserved functional genomic regions in Min6 cells (active promoters: 2370 genes, bivalent promoters: 1191 genes, inactive promoters: 3055 genes, active enhancers: 837 genes, poised enhancers: 648 genes, heterochromatin: 3308 genes). Results are displayed as mean of Log2(TPM+1) for each conserved functional genomic regions associated-genes. The violin plot displays median and quartile values for each group of genes.

This comparative analysis led us to consider the conservation level of the functional genomic regions characterized in Min6 cells (Figure 2) with the functional chromatin segments defined in mouse islets [21]. The functional genomic regions of Min6 cells were also conserved compared to functional chromatin segments defined in mouse islets (Figure 4B). Indeed, we showed that the most active promoters in Min6 cells were related to active promoter segments (segments A and B, Figure 4B, active promoter) while bivalent promoters were related to bivalent segments (segments L and M, Figure 4B, bivalent promoter) and inactive promoters were mostly related to silent segments (segments N to Q segments, Figure 4B, inactive promoter). Regarding the distal intergenic functional regions, the active enhancers mainly overlapped with the distal active segments (segments C to E, Figure 4B, active enhancers) whereas the poised enhancers were mainly related to the bivalent segments (K and M segments, Figure 4B, poised enhancers). Regarding the heterochromatin regions, these were mostly related to the inactive segments (mainly segment N, Figure 4B, heterochromatin). Each conserved functional genomic region was exemplified through a series of selected chromatin features (gene promoter or intergenic region, Figures 4C and 4D), such as the β-cell marker *Pdx1*, the α-cell marker somatostatin receptor 1 (*Sstr1*) or the ameloblast specific gene, Amelotin (*Amtn*). Finally, we selected the functional chromatin regions that were conserved between Min6 cells and mouse islets. To reach this aim, we selected only the functional chromatin regions of Min6 cells with more than 30% overlap with each related functional chromatin segment within the mouse islets. Subsequently, the expression level of the genes associated with these conserved genomic functional regions was analyzed (Figure 4E). The results notably showed that the genes associated with the conserved active chromatin regions (active promoter and active enhancers) are most expressed compared to the genes associated with the inactive regions in line with our data generated without applying the conservation filter (Figure 4E). Altogether, these results showed that the functional chromatin regions are mostly shared and conserved between Min6 cells and mouse islets, indicating that Min6 cells represent a pertinent β-cell model to explore the effect of HDAC inhibition at the genome-wide level.

### 3.4. HDAC inhibition differentially alters acetylation level of conserved functional genomic regions in Min6 cells

The correlation between the alteration of β-cell function and the increase of histone acetylation in response to HDAC inhibition (Figure 1) prompted us to interrogate the effects of TSA treatment on histone acetylation within previously characterized conserved functional genomic regions in Min6 cells. To reach this aim, Min6 cells were treated either with vehicle (DMSO 0.1%) or TSA (0.5 μM) during 16 hours and H3K9ac, H3K27ac and H3K27me3 were profiled through ChIP-seq experiments (Figure 5A). Compared to the vehicle condition, TSA treatment led to the detection of a higher number of peaks for both H3K9ac (27438 for TSA *versus* 13771 for DMSO) and H3K27ac (66347 for TSA *versus* 13451 for DMSO) (Figure 5B). Concomitantly, TSA treatment drastically reduced the number of peaks for H3K27me3 (17367 for DMSO *versus* 1253 for TSA) (Figure 5B). Consistent with the results from dot blot experiments (Figures 1A to D), these data however indicated that the increase of H3K9ac and H3K27ac signal intensity in response to TSA treatment may not be only related to an increase of signal intensity within constitutively acetylated regions but also to an increase in the number of *de novo* acetylated genomic sites. This was corroborated by the analysis of the genomic distribution of H3K9ac and H3K27ac in response to TSA treatment showing a significant redistribution of H3K9ac and H3K27ac enriched regions especially towards gene bodies and distal genomic regions for H3K9ac (Supplementar Figures 3A and B, H3K9ac) and gene bodies for H3K27ac (Supplementary Figures 3C and D, H3K27ac). The genomic distribution of H3K27me3 was less modified suggesting that it was rather the loss of signal than the genomic redistribution affecting H3K27me3 in response to TSA treatment (Supplementary Figures 3E and F, H3K27me3).

**Figure 5.**
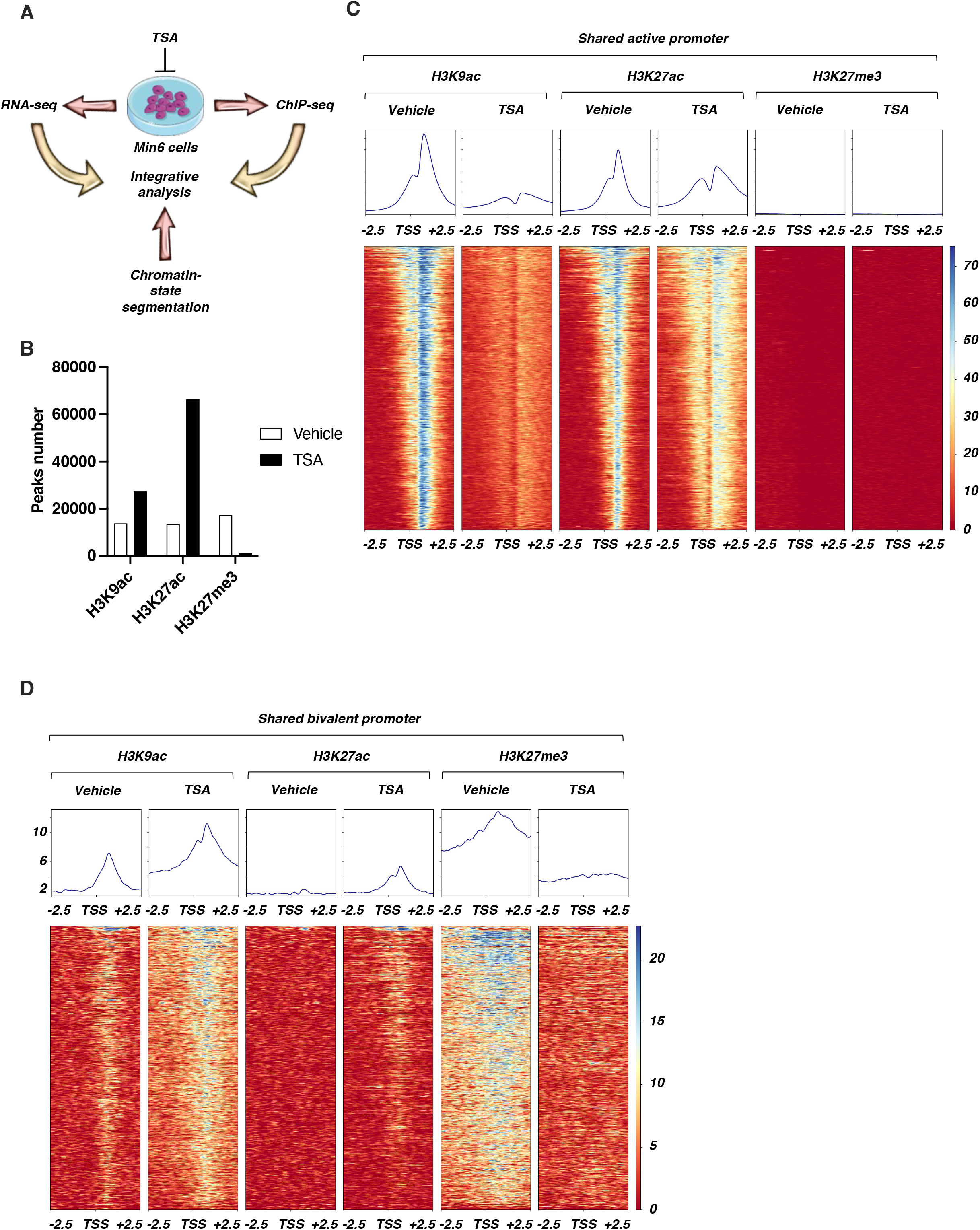
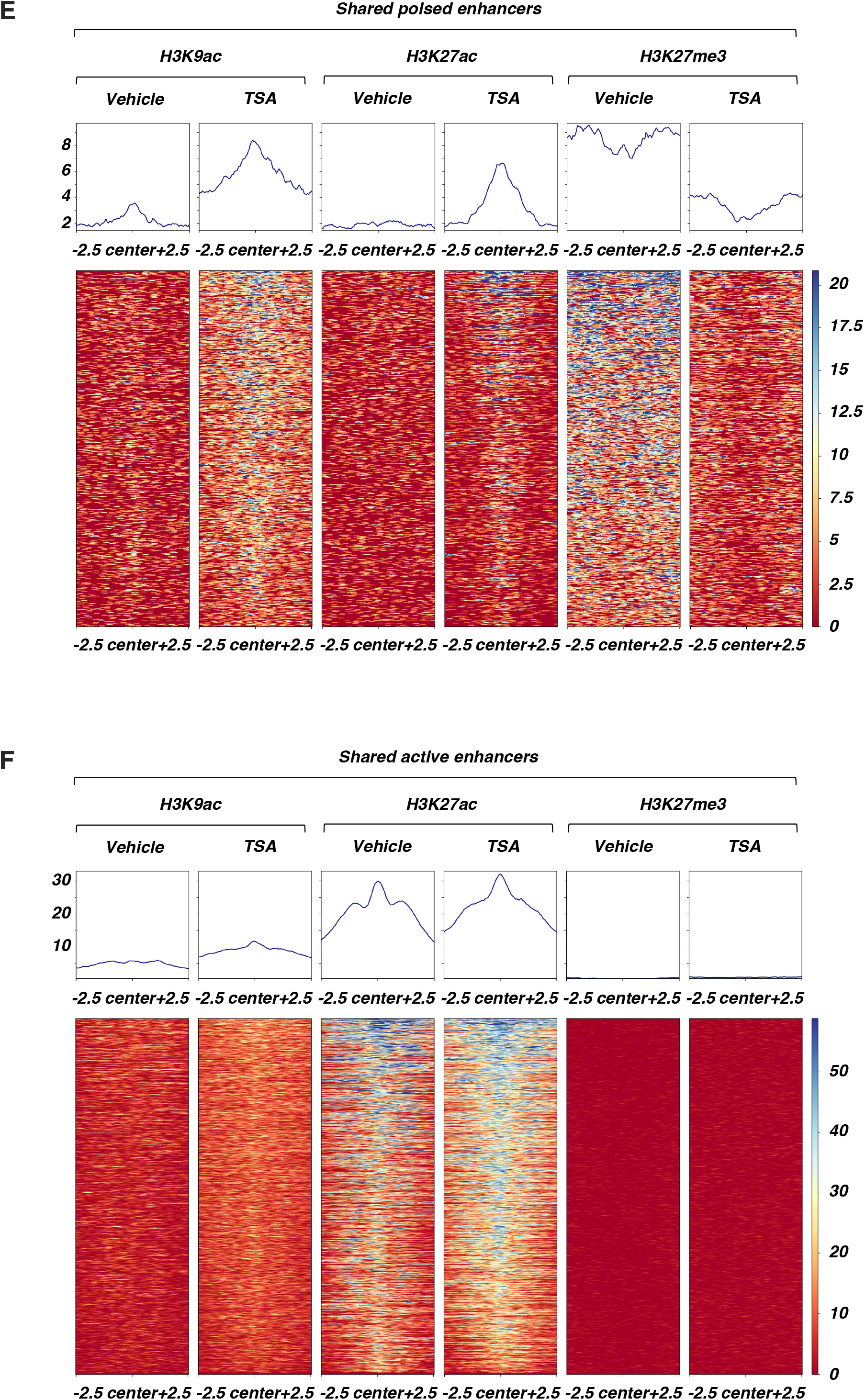
TSA treatment leads to a genomic redistribution of H3K9ac, H3K27ac and H3K27me3. **(A)** Scheme representing the strategy used in Min6 cells. **(B)** Number of H3K9ac, H3K27ac and H3K27me3 peaks in vehicle- and TSA-treated Min6 cells. **(C-F)** H3K9ac, H3K27ac and H3K27me3 signal in conserved active promoters (C), bivalent promoters (D), poised enhancers (E) and active enhancers (F) in vehicle- and TSA-treated Min6 cells. Heatmaps and mean signals centered on TSS +/- 2.5kb are represented.

Considering that a genomic redistribution of H3K9ac and H3K27ac occurred in response to TSA treatment, we then hypothesized that the levels of H3K9ac and H3K27ac acetylation could be altered within the conserved functional genomic regions described above (Figure 2 and Supplementary Figure 2). In addition, the H3K27me3 signal was also monitored in these regions. First, a promoter-focused analysis centered on the transcription start site (TSS) +/- 2.5 kb was performed. Surprisingly, this analysis revealed that H3K9ac signal was completely blunted after treating Min6 cells with TSA within shared active promoters while the H3K27ac signal was weakly affected or even slightly increased in the close vicinity of the promoters whereas H3K27me3 signal was not modulated (Figure 5C). Conversely, an increase of H3K9ac and, to a lesser extent, H3K27ac signal, associated with a decrease of H3K27me3, was detected within shared bivalent promoters upon TSA condition (Figure 5D). Considering shared inactive promoters, neither H3K9ac, H3K27ac nor H3K27me3 signal was modulated by TSA treatment (Supplementary Figure 4A). This promoter-based analysis at genome-wide level showed that TSA treatment led to distinct effects on histone acetylation depending on the basal activation level of the promoters. Acetylation level in response to TSA treatment was next interrogated within conserved functional distal intergenic regions at genome-wide level focusing on the center of the region +/- 2.5 kb. Within shared poised enhancers, both H3K9ac and H3K27ac signal was increased whereas H3K27me3 signal was decreased (Figure 5E) while shared active enhancers were weakly enriched with H3K9ac and H3K27ac without modulation of H3K27me3 signal upon TSA treatment (Figure 5F). Finally, shared heterochromatin-focused analysis showed that H3K27me3 was decreased within these regions whereas H3K9ac and H3K27ac signals were weakly affected (Supplementary Figure 4B). Taken together, these genome-wide analyses suggest that TSA treatment in Min6 cells differentially impacts acetylation profile depending on the type of functional genomic regions. This implied that gene expression level could be directly affected by these changes after genomic reprogramming of acetylation.

### 3.5. Genomic redistribution of histone acetylation upon HDAC inhibition directly reprograms the transcriptome in Min6 cells

We interrogated the functional outcomes of the modulation of histone acetylation profile within conserved functional genomic regions through transcriptome-wide analysis in Min6 cells. To reach this aim, RNA-seq was then performed on 16 h vehicle- and TSA-treated Min6 cells. By applying an adjusted p-value cut-off threshold at 0.05 (Padj<0.05), 5636 genes were significantly downregulated and 6571 genes were significantly upregulated (Supplementary Table S3). Given this high number of deregulated genes, we applied a second cut-off threshold based on Log2 fold change (Log2FC) to select only the most deregulated genes. Using these two cut-off thresholds, 2626 genes were downregulated (Padj<0.05, Log2FC<-1) and 3877 genes were upregulated (Padj<0.05, Log2FC>1) upon TSA treatment (Figure 6A, Supplementary Table S3). Ingenuity pathway analysis (IPA) revealed that downregulated genes were associated with canonical pathways involved in kinetochore metaphase signaling (-Log10(P)=16), mitotic roles of Polo-like kinase (-Log10(P)=9), but most interestingly, with insulin secretion (-Log10(P)=9), Maturity Onset of Diabetes of Young (MODY, -Log10(P)=7) and GPCR-mediated nutrient sensing (-Log10(P)=5, Figure 6B, Supplementary Table S4). In line with this, several key β-cell genes were negatively affected by TSA treatment, such as *Slc2a2, Nkx2-2, G6pc2, Kcnj11, Mafa, Pdx1* or *P2ry1* (Supplementary Figure 5A and Supplementary Table S3). Conversely, upregulated genes were associated to pathways unrelated to β-cell functions, such as axonal guidance, pulmonary fibrosis or cardiac hypertrophy (Figure 6C, Supplementary Table S5). Based on these results, the deregulated genes were classified according to the conserved functional genomic region with which they were associated and the proportion of each group was plotted in a pie chart. Among the downregulated genes, this analysis showed that more than 25% were genes with an active promoter (Figure 6D) and 10% of the downregulated genes were associated with heterochromatin regions. Genes associated with other functional genomic regions were marginally represented (less than 15%). In addition, this analysis showed that 48% were not associated with any previously characterized functional genomic region, likely due to a lower analytical power related to the restricted number of histone marks profiled in this study. These results were directly correlated to the functional genomic data showing that the active promoters harbored a drastic decrease in the H3K9ac signal potentially responsible to gene expression downregulation (Figure 5B).

**Figure 6.**
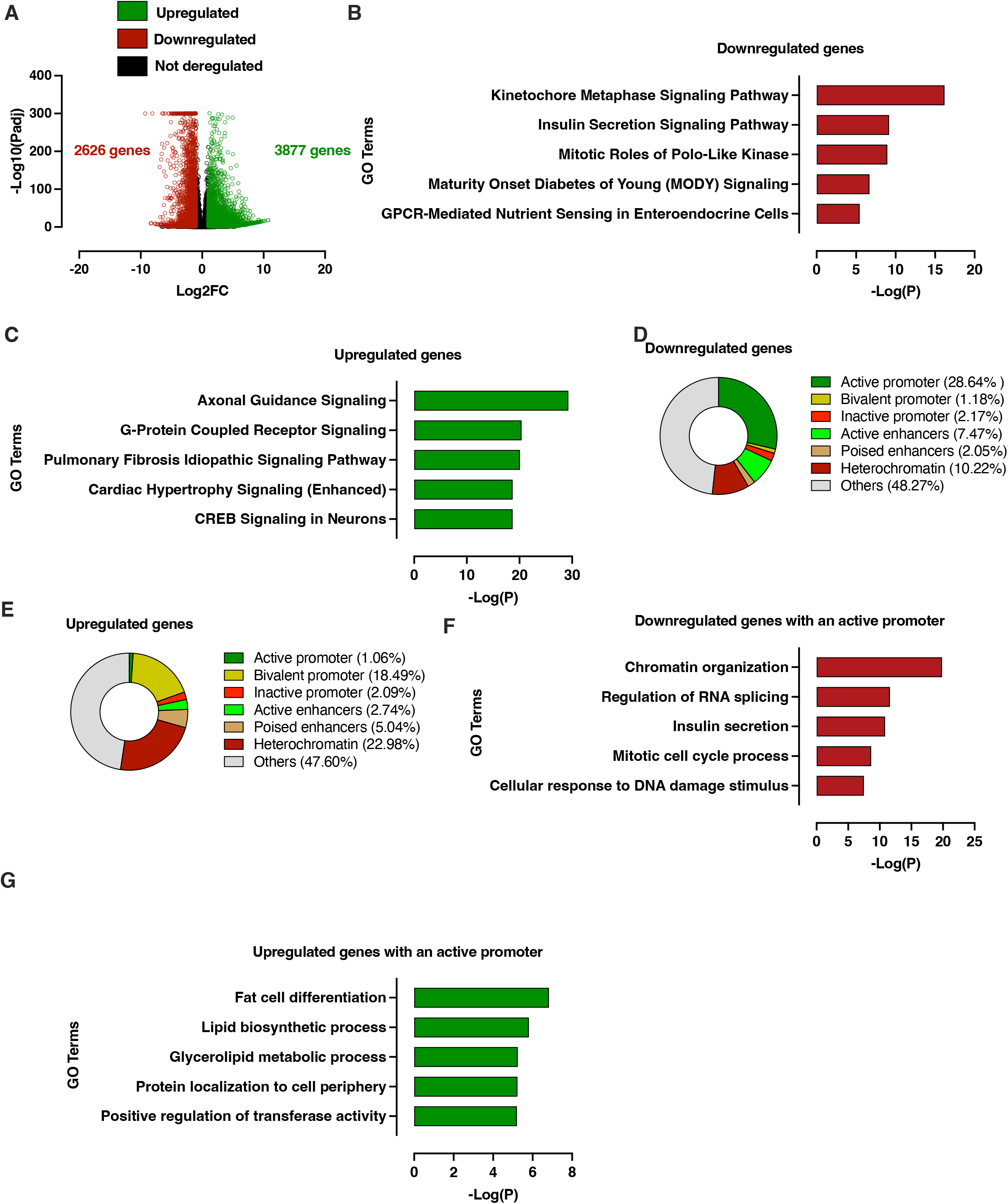
Redistribution of histone acetylation at genome-wide level upon TSA treatment in Min6 cells is associated with differential gene expression at transcriptome-wide level. **(A)** Volcano plot displaying downregulated (red circles), upregulated (green circles) and not deregulated genes (black circles) in TSA-treated cells compared to vehicle-treated cells according to two cut-off thresholds based on adjusted p-value (Padj) and Log2 fold change (Log2FC). **(B-C)** Ingenuity Pathway analysis (IPA) of downregulated (B) and upregulated (C) genes in Min6 cells upon TSA treatment. **(D-E)** Pie chart displaying proportion of TSA-dependent downregulated (D) and upregulated (E) genes in Min6 cells associated with a specific functional genomic region. **(F-G)** Metascape analysis of genes with an active promoter that are downregulated (F) and upregulated (G) upon TSA treatment.

Regarding upregulated genes, 46% were associated with silent functional genomic regions (bivalent promoter, 18%, heterochromatin, 23% and poised enhancers, 5%) while genes associated with active functional genomic regions were less represented (active promoter, 1% and active enhancers, 3%) (Figure 6E). Again, 48% of the downregulated genes were not associated with previously defined functional genomic regions. This analysis however highlighted the direct link between altered epigenomic profile and increased gene expression in response to TSA treatment since previous data demonstrated that silent regions showed either an enrichment in H3K9ac/H3K27ac associated with a depletion in H3K27me3 (*i.e*., bivalent promoter and poised enhancers; Figures 5D and 5E, respectively) or only depletion in H3K27me3 (*i.e*., heterochromatin; Supplementary Figure 4B). These observations were exemplified through a series of selected β-cell (*Pdx1* and *Mafa*) and α-cell markers (*Arx* and *Mafb*) demonstrating that β-cell genes were depleted in H3K9ac/H3K27ac marks, whereas α-cell genes were slightly increased in H3K9ac/H3K27ac marks but depleted in H3K27me3 mark (Supplementary Figure 5C).

Consistent with the analysis of the effects of TSA on H3K9ac, H3K27ac and H3K27me3 levels within the conserved functional genomic regions in Min6 cells, these results showed a direct functional repercussion of the epigenomic alteration of these regions on the regulation of gene expression. This was all the more likely since the genes associated with inactive promoters whose H3K9ac, H3K27ac and H3K27me3 profile were marginally altered in response to TSA treatment were not enriched neither in downregulated genes (2.17%, Figure 6D) nor in upregulated genes (2.09%, Figure 6E). Gene ontology analysis using Metascape revealed that most of the downregulated genes upon TSA treatment were associated to chromatin organization, RNA splicing and insulin secretion (Figure 6F). Conversely, upregulated genes were enriched in pathways that are unrelated to β-cell function, such as fat cell differentiation or lipid biosynthetic process (Figure 6G). Altogether, these data suggest that TSA treatment functionally affects cell fate through an epigenome-wide remodeling associated to transcriptome changes that alter β-cell identity and function.

### 3.6. The transcriptome of FACS-sorted TSA-treated mouse β cells identifies partially conserved genomic mode of action of TSA-mediated HDAC inhibition

Having demonstrated a direct link between the epigenome alteration and modulation of gene expression in Min6 cells in response to HDAC inhibition, we next sought to determine whether the transcriptome as well as the associated-chromatin features could be conserved in mouse β cells. Pancreatic islets of Langerhans isolated from RIPCre-tdTomato mice expressing the red fluorescent protein td-Tomato under the control of the β-cell specific Cre recombinase were treated by either vehicle (DMSO 0.1%) or TSA (0.5 μM, 16 h) and then subjected to FACS analysis to sort Tomato positive cells, corresponding to a β-cell enriched population. RNA-seq experiment was performed on this β-cell enriched population (Figure 7A). In order to compare these data with those obtained in Min6 cells, an adjusted p-value and Log2FC cut-off thresholds (Padj<0.05 and -1>Log2FC>1) were applied, respectively. In these conditions, 1182 genes were significantly downregulated and 1331 genes were significantly upregulated upon TSA treatment (Figure 7B, Supplementary Table S6). IPA further confirmed that downregulated genes from TSA-treated sorted β-cells were mostly associated to insulin secretion and MODY signaling pathways (Figure 7C, Supplementary Table S7), whereas upregulated genes were found to be involved in the regulation of axonal guidance signaling, molecular mechanisms of cancer or WNT/ β-catenin signaling (Figure 7D, Supplementary Table S8).

**Figure 7.**
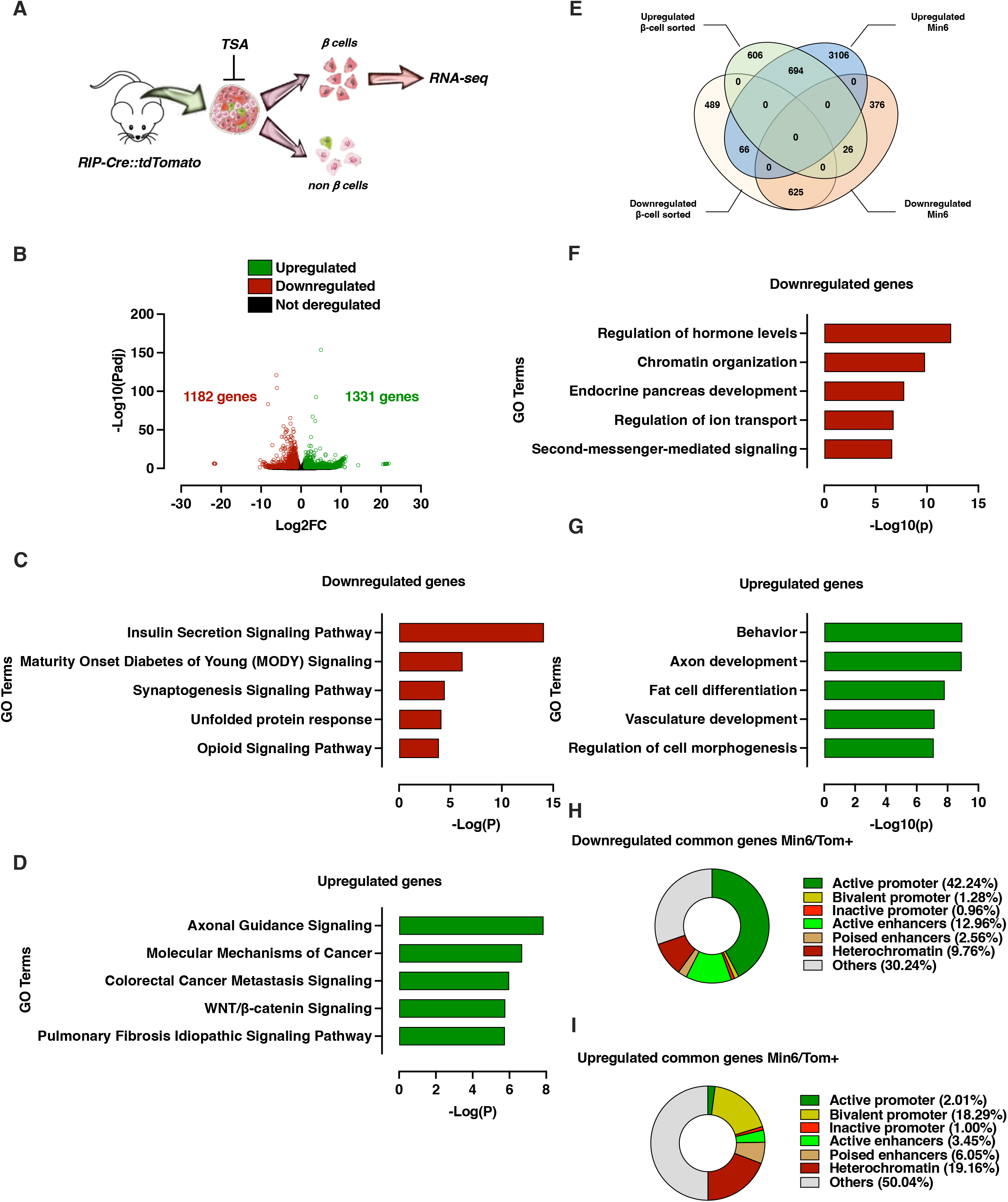
Transcriptome of FACS-sorted β-cell from *ex vivo* TSA-treated mouse Langerhans islets is partly shared with Min6 cells. **(A)** Schematic representation of β-cell preparation from mouse Langerhans islets. **(B)** Volcano plot displaying downregulated (red circles), upregulated (green circles) and not deregulated genes (black circles) in β-cell from TSA-treated mouse Langerhans islets compared to β-cell from vehicle-treated mouse Langerhans islets according to two cut-off thresholds based on adjusted p-value (Padj) and Log2 fold change (Log2FC). **(C-D)** Top 5 canonical pathways identified by IPA of downregulated (C) and upregulated (D) genes in TSA-treated mouse sorted-β cells. **(E)** Venn diagram identifying common TSA-dependent up- and downregulated genes between Min6 cells and β-cell from mouse Langerhans islets. **(F-G)** GO analysis of common TSA-dependent downregulated (F) and upregulated (G) genes. Gene ontology analysis was performed using Metascape by filtering output only on GO Biological process. The 5 more enriched GO Biological processes are displayed as -Log10(q) (*i.e*., -Log10(Padj)). **(H-I)** Pie chart displaying proportion of TSA-dependent downregulated (H) and upregulated (I) genes in β-cell from mouse Langerhans islets associated with a specific functional genomic region.

In order to focus our analysis on conserved transcriptomic pattern between FACS-sorted β cells and Min6 cells, we next intersected the related dataset in a Venn diagram (Figure 7E). Among upregulated genes defined in FACS-sorted TSA-treated β-cells, 53% (694 genes) and 2% (26 genes) were shared with upregulated and downregulated genes detected in Min6 cells, respectively. Of note, 46% of upregulated genes were specific to FACS-sorted TSA-treated β-cell. Downregulated genes in TSA-treated β-cell sorted were also mostly conserved compared to downregulated genes in TSA-treated Min6 cells since 53% were shared whereas only 5.6% were upregulated in Min6 cells. TSA treatment also induced a β-cell sorted specific transcriptomic signature since 41% of genes were not overlapping neither with downregulated nor upregulated genes in Min6 cells. This analysis suggested that HDAC inhibition through TSA treatment induced a transcriptomic signature resulting at least partly from conserved molecular mechanisms between *ex vivo* TSA-treated β-cell and the Min6 β-cell model.

To go further in the functional characterization of these conserved deregulated genes upon TSA treatment, a gene ontology analysis using IPA was conducted on both conserved down- and upregulated genes. Conserved downregulated genes were associated with biological processes mostly related to β-cell function (*e.g*., regulation of hormone levels and endocrine pancreas development, Figure 7F) whereas conserved upregulated genes were rather associated with biological processes unrelated to β-cell function (*e.g*., axon development and fat cell differentiation, Figure 7G). Altogether, this analysis suggested that the alteration of β-cell function upon HDAC inhibition could be conserved and related to selective downregulation of β-cell specific genes. To ascertain this hypothesis, we conducted a Gene Set Enrichment Analysis (GSEA) to monitored β-cell specific gene set enrichment within conserved downregulated genes (Supplementary Figure 5B). As expected, β-cell specific genes were mostly enriched in vehicle-compared to TSA-treated β-cell-sorted, thus supporting our gene ontology analysis.

With the aim to connect our transcriptomic data with functional genomic data from mouse β cells, we next sought to determine the functional genomic regions to which conserved deregulated genes were associated. By using the previously defined conserved functional genomic regions (Figure 2B), we were able to demonstrate that most of the conserved downregulated genes were associated to active regions (active promoter, 42% and active enhancers, 13%) (Figure 7H) whereas upregulated genes were rather associated to silent regions (bivalent promoter, 18%, poised enhancers, 6% and heterochromatin, 19%) (Figure 7I). Of note, these results were close to results obtained in Min6 cells (Figures 6D and 6E) suggesting that a part of genomic regions functionally affected upon TSA treatment were conserved between β-cell-sorted and Min6 cells.

### 3.7. Transcriptome-wide analysis of EndoC-βH1 and human islets upon HDAC inhibition identifies conserved TSA-sensitive genes related to β-cell identity

Given that HDAC inhibition led to an important redistribution of histone acetylation at genome-wide level leading to a drastic modulation of gene expression at transcriptome-wide level both in Min6 and mouse β-cell sorted cells, we next asked whether TSA treatment could also reprogram the transcriptome of human β cells. Consequently, EndoC-βH1 cells and human pancreatic islets were independently treated either with vehicle (DMSO 0.1%) or TSA (0.5 μM) during 16 hours and the resulting transcriptome was recorded through RNA-seq experiments as depicted in Figures 8A and 8B, respectively.

**Figure 8.**
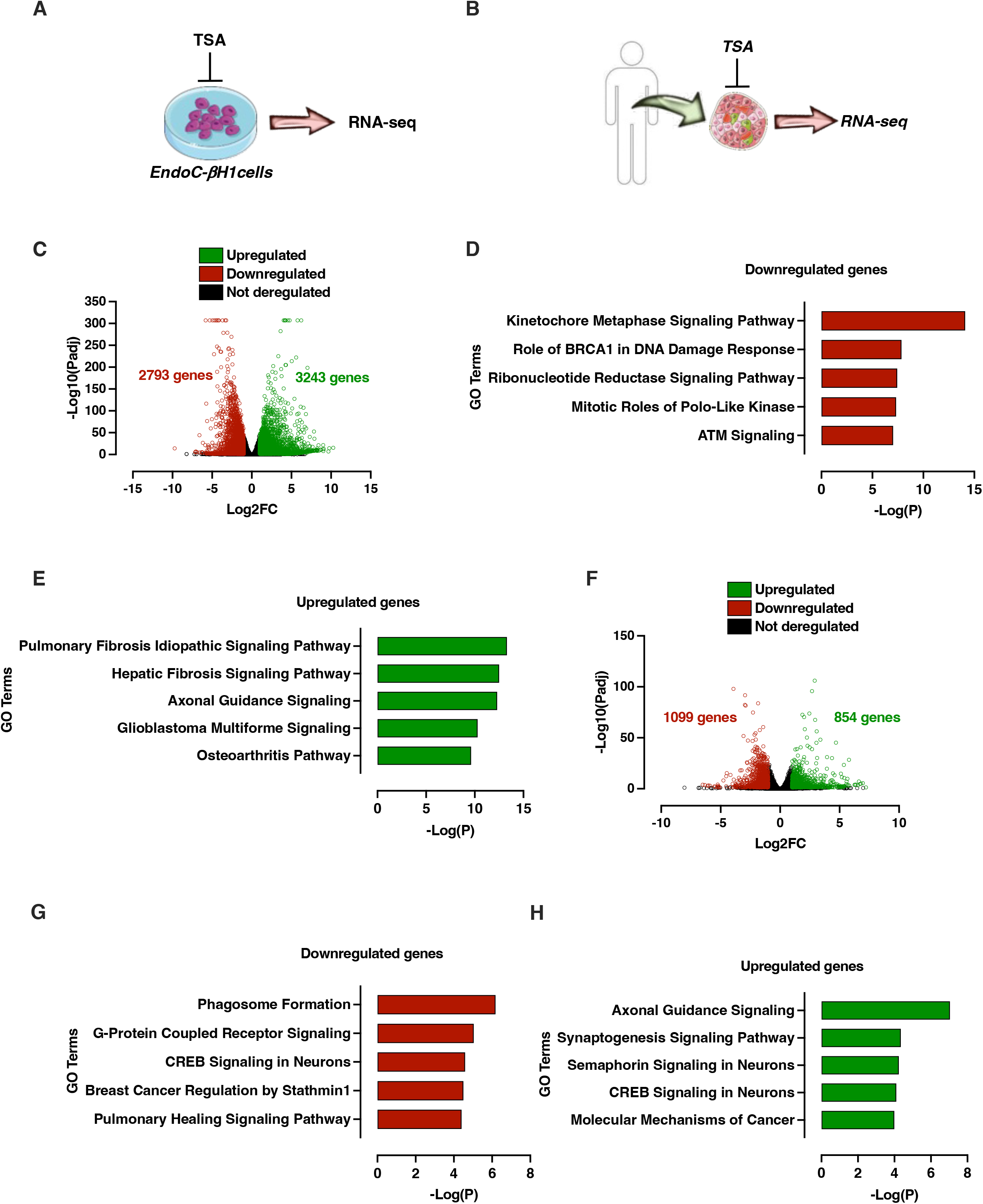
**(A-B)** Schematic representation of EndoC-βH1 and human islets treatment by TSA and subsequent RNA-seq analysis. **(C)** Volcano plot displaying downregulated (red circles), upregulated (green circles) and not deregulated genes (black circles) in EndoC-βH1 control and TSA-treated cells. **(D-E)** Top 5 canonical pathways identified by IPA of downregulated (D) and upregulated (E) genes in TSA-treated EndoC-βH1 cells. **(F)** Volcano plot displaying downregulated (red circles), upregulated (green circles) and not deregulated genes (black circles) in TSA-treated human islet compared to untreated controls. **(G-H)** Top 5 canonical pathways identified by IPA of downregulated (D) and upregulated (E) genes in TSA-treated human pancreatic islets.

In EndoC-βH1 cells, by applying an adjusted p-value cut-off threshold at 0.05 (Padj<0.05), 4717 genes were significantly downregulated and 5443 genes were significantly upregulated (Supplementary Table S9). As in Min6 cells, we applied a second cut-off threshold based on Log2 fold change (Log2FC) to select only the most deregulated genes. Using these two cut-off thresholds, 2793 genes were downregulated (Padj<0.05, Log2FC<-1), 3243 genes were upregulated (Padj<0.05, Log2FC>1) upon TSA treatment (Figure 8C and Supplementary Table S9). These results showed that the transcriptome of EndoC-βH1 cell line was highly sensitive to and profoundly affected by TSA treatment, as observed in the Min6 cell line and FACS-sorted β cells. Pathway analysis of downregulated genes suggested that TSA affects pathways controlling cell cycle and cell proliferation, but also insulin secretion (Log(pVal)= 1.8; Z-score=-5.38; Figure 8D, Supplementary Tables S9 and S10). Amongst genes that are downregulated and identified as β-cell identity genes [22], *ABCC8, MAFA, GLP1R* and *PDX1* were found to be negatively affected. Conversely, TSA-mediated upregulated genes were associated to canonical pathways that were unrelated to β-cell functions, such as fibrosis, axonal guidance or osteoarthritis pathways (Figure 8E, Supplementary Tables S9 and S11). In non-diabetic human islets, 1099 genes were significantly downregulated and 854 genes were significantly upregulated, after applying cut-off thresholds (Padj<0.05, -1>Log2FC>1, Figure 8F, Supplementary Tables S12). Among the canonical pathways that were affected in downregulated genes, phagosome formation, G-protein coupled receptor signaling and CREB signaling in neurons were the most represented (Figure 8G and Supplementary Table S13). Again, insulin secretion signaling pathway was found to be affected by TSA treatment (Log(pVal)= 1.64; Z-score=-3.15; Supplementary Table S13). For upregulated genes, IPA revealed that genes involved in pathways unrelated to pancreatic islet function were enriched following TSA treatment, such as axonal guidance signaling, synaptogenesis or semaphorin signaling pathways (Figure 8H and Supplementary Table S14). Although we did not measure the GSIS of EndoC-βH1 and human islets in response to TSA treatment, these results suggest that TSA induces a profound transcriptomic reprogramming in human β-cell line.

## 4. Discussion

Here we show that blocking HDAC activity with TSA profoundly impairs insulin secretion. This functional effect is associated to an epigenomic reprogramming, associated to loss of β-cell identity genes, in mouse as well as in human β cells or islets. Although the contributions of HDACs to endocrine pancreas development are well circumscribed [23; 24], our results demonstrate that they also contribute to the maintenance of mature β-cell identity, plasticity and function.

It has been recently suggested that enhancement of ROS (reactive oxygen species) production upon TSA treatment in Min6 cells could have negative effects of HDACi on β-cell function [25]. An increase of IRS2 (Insulin Receptor Substrate) expression in response to TSA treatment in Min6 cells has been reported to be causal of the impairment of β-cell function upon HDAC inhibition [26]. Our transcriptomic analyses reveal that TSA treatment decreases rather than increases IRS2 expression (Supplementary Table S3) and mostly affects the expression of a large set of genes involved in various biological process, suggesting that the effect of HDACi on β-cell function is probably more complex than anticipated. Indeed, among deregulated genes in response to TSA treatment, most of β-cell specific genes are downregulated whereas a series of α-cell specific genes as well as genes unrelated to endocrine functions are upregulated. In accordance with previous results from Kubicek and colleagues observed in TSA-treated murine β-TC6 cells [15], these results suggest that global HDAC inhibition leads to an alternative transcriptional program leading to a switch from β-cell identity towards an undefined alternative identity partly overlapping with α-cell identity. Also in line with the analysis restricted to some β-cell markers from Yamoto E [25], our data confirm that β-cell identity is effectively directly impaired by HDAC inhibition.

In this context, we demonstrate here that this alternative transcriptional program is likely consecutive to a drastic alteration of epigenome upon HDAC inhibition in β-cell, especially through modification of histone acetylation profile within promoters. Whereas the definition of H3K9ac-enriched promoters at genome-wide level is consistent with global increase of histone acetylation level, it is much more counterintuitive to identify H3K9ac-depleted promoters upon HDAC inhibition. Indeed, we found most of the β-cell specific genes promoters surprisingly H3K9ac-depeleted in response to HDAC inhibition and correlated to a decrease of gene expression. As TSA is an unspecific class I and class II HDAC inhibitor, this experimental evidence suggests that one or several class I and/or class II HDAC enzymes could play a direct active role in the regulation of expression of genes involved in the β-cell identity. In accordance with this hypothesis, class I (*i.e*., HDAC1, 2 and 3) and class IIb (*i.e*., HDAC6) HDAC enzymes were shown to be bound within promoter of transcriptionally active gene promoters in human CD4+ T cells [27]. However, we did not demonstrate any involvement of neither HDAC1 nor HDAC6 in the direct regulation of transcriptionally active genes in β cell (data not shown) suggesting that (*i*) conversely to observations from CD4+ T cells, HDAC1 and HDAC6 are not directly involved in the regulation of β-cell specific genes, (*ii*) other HDAC could be implicated in this process such as HDAC2 and/or HDAC3 [13; 27]. Alternatively, depletion of acetylation in promoter of transcriptionally active genes upon HDAC inhibition in β cell could be consecutive to the loss of a histone acetyl transferase enzyme binding within promoter of these genes which has been recently proposed for CBP/p300 as an alternative mechanistic model explaining promoter histone deacetylation in response to HDAC inhibition in endothelial cells [28].

We also provided evidences that HDAC inhibition also leads to enhancer remodeling. Enhancers are cell specific non-coding elements of the genome involved in long-distance cell-specific regulation of gene expression directly related to cell identity, especially β-cell identity [29]. It has been recently reported that the HDAC inhibition using largazole induces a H3K27ac-depletion at enhancers in HCT116 cells [30] suggesting a role of HDAC in enhancer activity. Although it remains to experimentally demonstrate that HDACs directly bind to β-cell enhancers, our results emphasize a crucial role of HDAC enzymes in enhancer activity to maintain an appropriate transcriptional pattern. At a mechanistic level, the level of enhancer activity is especially directly related to the amount of local production of enhancers RNA (eRNA) [31] playing a direct role in the regulation of enhancer target genes expression [32] In addition, as eRNA are transcribed from active enhancers, they exhibit tissue and lineage specificity, and serve as markers of cell state and function [33]. Consequently, we hypothesize that remodeling of β-cell enhancers upon HDAC inhibition could lead to an alteration of the β-cell specific eRNA expression profile, thus partly contributing to affect specific gene expression profile as recently pointed out in human BT474 cells [34]. To date, eRNAs are not well characterized in β cells and the link between HDAC and eRNA needs to be further investigated to better define β-cell identity as well as improve molecular knowledge on β-cell function. It would be interesting to investigate the role of distinct HDAC using selective or specific HDACi [35].

Although this study connects the negative effects of HDACi on β-cell function to the global remodeling of epigenome (*i.e*., histone acetylation) at genome-wide level, we cannot rule out that alteration of acetylome at the proteome-wide level upon HDAC inhibition is related to the observed phenotype. Indeed, acetylation is a largely widespread conserved post-translational modification also affecting non-histone proteins/enzymes to regulate their functions [36]. As acetylome recently defined in rat islet is enriched in metabolic pathways of β cells related to nutrient sensing [37], we propose that HDAC inhibition using hydroxamic acid such as TSA leads to a global protein deacetylation involved in these pathways impairing protein/enzyme functions concomitantly to epigenome remodeling inducing in turn β-cell failure. Nevertheless, given the specificity of action of TSA on class I and II HDAC, this implies that these two classes of enzymes play a role on both nuclear and cytoplasmic non-histone protein deacetylation process, which has not been yet reported in β cell whereas Sirt3 belonging to class III HDAC takes part to this process in rat Ins1 cells [37].

In summary, based on a pharmacological approach using the pan-HDAC inhibitor TSA, we provide new information indicating that HDAC inhibition negatively acts on β-cell transcriptional program through epigenome-wide remodeling, leading to an alteration of β-cell functional properties. These results also indicate that β-cell identity is likely under the control of HDAC activity acting both on gene expression as well as enhancer activity. Other investigations are currently in progress to better define the role of each HDAC in this process.

## Funding

This work was supported by the Agence Nationale de la Recherche (ANR) grants (EGID ANR-10-LABX-46; LIGAN-PM Equipex 2010 ANR-10-EQPX-07-01; ANR-16-IDEX-0004 ULNE; ANR BETAPLASTICITY ANR-17-CE14-0034), EFSD, INSERM, CNRS, Université de Lille, Institut Pasteur de Lille (CTRL Melodie), Fondation pour la Recherche Médicale (FDT202106013015, EQU202103012732), I-SITE ULNE (EpiRNAdiab Sustain grant), Conseil Régional Hauts de France, Métropole Européenne de Lille, Société d’Accélération du Transfert de Technologie Nord and Société Francophone du Diabète.

## Acknowledgments

The authors thank the members of the INSERM U1283-EGID and INSERM UMR1167-RID-AGE for helpful discussions. We are grateful to Yannick Campion for his support and his administrative contribution during this project. Human islets were provided through the JDRF award 31-2008-416 (ECIT Islet for Basic Research program). The authors thank “France Génomique” consortium (ANR-10-INBS-009), UMS2014-US41 and the Experimental Resources platform from Université de Lille.

## Authors contributions

Conceptualization, F.O., and J.S.A.; Methodology, F.O., M.M., Am.B. and J.S.A.; Investigation, F.O., M.M., M.D., B.T., L.B., C.B., E.D., S.A., Al.B., E.B., O.M.C., L.P., G.P., L.R., C.C., F.B. and E.C.; Resources, C.G., J.E., D.D., J.K.C., F.P., B.S., P.F., and Am.B.; Data curation, F.O., M.M, M.D., L.B., S.A., and Al.B.; Writing – original draft, F.O., M.M., and J.S.A.; Writing – Review & Editing, J.E., P.F., and Am.B.; Visualization, F.O., M.M., and J.S.A.; Supervision, F.O., Am.B. and J.S.A.; Funding acquisition, J.S.A.

## Conflict of interest

The authors declare that there is no conflict of interests regarding the publication of this article.

## Notes

### Competing Interest Statement

The authors have declared no competing interest.

