## Supplementary material for "The HDAC inhibitor trichostatin A impairs pancreatic β-cell function through an epigenome-wide reprogramming": supp_text

**Footnotes**

[†] Co-senior authors

[*] Corresponding authors:

Running title: HDAC and β-cell function

**Supplementary Table S1.** List of antibodies used for Dot Blot or ChIP

| Product Name | Species | Provider | Product Reference | Dilution/quantity |
|---|---|---|---|---|
| Anti-IgG antibody | Mouse | Santa Cruz Biotechnologies | #sc2025 | WB : 1/1000 |
| Anti-H3K4me3 antibody | Mouse | Active motif | #39159 | ChIP : 2 µg/IP |
| Anti-H3K27me3 antibody | Mouse | Active motif | #61017 | Dot blot : 1/1000<br>ChIP : 2 µg/IP |
| Anti-H3K27ac antibody | Mouse | Active motif | #39685 | Dot blot : 1/1000<br>ChIP : 2 µg/IP |
| Anti-H3K9ac antibody | Rabbit | Abcam | #ab4441 | Dot blot : 1/1000<br>ChIP : 2 µg/IP |
| Anti-H3K4me1 antibody | Rabbit | Active motif | #39297 | ChIP : 2 µg/IP |
| Anti-H3-pan-acetylated antibody | Rabbit | Abcam | #ab47915 | Dot blot : 1/1000 |
| Anti-H3 antibody | Rabbit | Abcam | #ab1791 | Dot blot : 1/1000 |
| Anti-Mouse HRP antibody | Rabbit | Sigma | A9044 | 1/10000 |
| Anti-Rabbit HRP antibody | Goat | Sigma | A9169 | 1/10000 |

**Supplementary Table S2.** List of oligonucleotides used in qPCR experiments.

| Official symbol | Species | Sequence | Forward / Reverse |
|---|---|---|---|
| *CYPA (Cyclo)* | Mouse | ATGGCACTGGCGGCAGGTCC | Forward |
| | | TTGCCATTCCTGGACCCAAA | Reverse |
| *Pdx1* | Mouse | ATTGTGCGGTGACCTCGGGC | Forward |
| | | GATGCTGGAGGGCTGTGGCG | Reverse |
| *MafA* | Mouse | TCCGACTGAAACAGAAGCGG | Forward |
| | | CTCTGGAGCTGGCACTTCTC | Reverse |
| *Foxo1* | Mouse | TGCCCAACCAAAGCTTCCCACA | Forward |
| | | TGGACTGCTCCTCAGTTCCTGCT | Reverse |
| *Ins1* | Mouse | GCCAAACAGCAAAGTCCAGG | Forward |
| | | GTTGAAACAATGACCTGCTTGC | Reverse |

**Supplementary Table S3.** List of differentially expressed genes in Min6 cells treated with TSA.

**Supplementary Table S4.** Ingenuity pathway analysis of down-regulated genes in Min6 cells treated with TSA.

**Supplementary Table S5.** Ingenuity pathway analysis of up-regulated genes in Min6 cells treated with TSA.

**Supplementary Table S6.** List of differentially expressed genes in sorted beta cells treated with TSA.

**Supplementary Table S7.** Ingenuity pathway analysis of down-regulated genes in sorted beta cells treated with TSA.

**Supplementary Table S8.** Ingenuity pathway analysis of up-regulated genes in sorted beta cells treated with TSA.

**Supplementary Table S9.** List of differentially expressed genes in EndoCBH1 cells treated with TSA.

**Supplementary Table S10.** Ingenuity pathway analysis of down-regulated genes in EndoCBH1 cells treated with TSA.

**Supplementary Table S11.** Ingenuity pathway analysis of up-regulated genes in EndoCBH1 cells treated with TSA.

**Supplementary Table S12.** List of differentially expressed genes in pancreatic human islets treated with TSA.

**Supplementary Table S13.** Ingenuity pathway analysis of down-regulated genes in pancreatic human islets treated with TSA.

**Supplementary Table S14.** Ingenuity pathway analysis of up-regulated genes in pancreatic human islets treated with TSA.

**Supplementary Figure 1. (A-B)** Annexin V (A) and propidium iodide (B) labelling of TSA-treated Min6 cells (0.5 µM, 16h, n=3). Vehicle (DMSO 0.1%, n=3) was used as negative control. Annexin - : annexin V negative cells, Annexin + : annexin V positive cells. IP - : propidium iodide negative cells, IP + : propidium iodide positive cells. Results in A and B are displayed as % of single cells recorded by FACS +/- SEM. * $p<0.05$, ns: not significant.

**Supplementary Figure 2. (A-D)** Individual genomic distribution of H3K4me3, H3K4me1, H3K27ac and H3K27me3 in Min6 cells. The percentage of peaks for each histone mark within 4 distinct genomic segments is displayed in a pie chart (promoter (TSS +/- 2.5 kb), gene body, downstream of genes and distal intergenic regions).

**Supplementary Figure 3. (A-F)** Individual genomic distribution of H3K9ac (A-B), H3K27ac (C-D) and H3K27me3 (E-F) in vehicle- (A, C and E) and TSA-treated (B, D and F) Min6 cells. The percentage of peaks for each histone mark within 4 distinct genomic segments is displayed in a pie chart (promoter (TSS +/- 2.5 kb), gene body, downstream of genes and distal intergenic regions.

**Supplementary Figure 4. (A)** H3K9ac, H3K27ac and H3K27me3 signal in conserved inactive promoters in vehicle- and TSA-treated Min6 cells. Heatmap and mean signal centered on TSS +/- 2.5kb are displayed. **(B)** H3K9ac, H3K27ac and H3K27me3 signal in conserved heterochromatin in vehicle- and TSA-treated Min6 cells. Heatmap and mean signal centered on TSS +/- 2.5kb are displayed.

**Supplementary Figure 5. (A)** Heatmap representing gene expression levels of β-cell genes in vehicle and TSA-treated Min6 cells. **(B)** Enrichment plot from Gene Set Enrichment Analysis (GSEA). GSEA was conducted with 60 genes set enriched in β cells. **(C)** Examples of functional genomic regions of β-cell (*Pdx1, Mafa)* and α-cell genes (*Arx, Mafb*) in vehicle and TSA-treated Min6 cells showing H3K9ac, H3K27ac and H3K27me3 marks. Figures are adapted from the IGB genome browser screenshots.
