## Supplementary figures and images for "The HDAC inhibitor trichostatin A impairs pancreatic β-cell function through an epigenome-wide reprogramming"

### Supp_figures

Oger *et al.*, Supplementary Figure 1

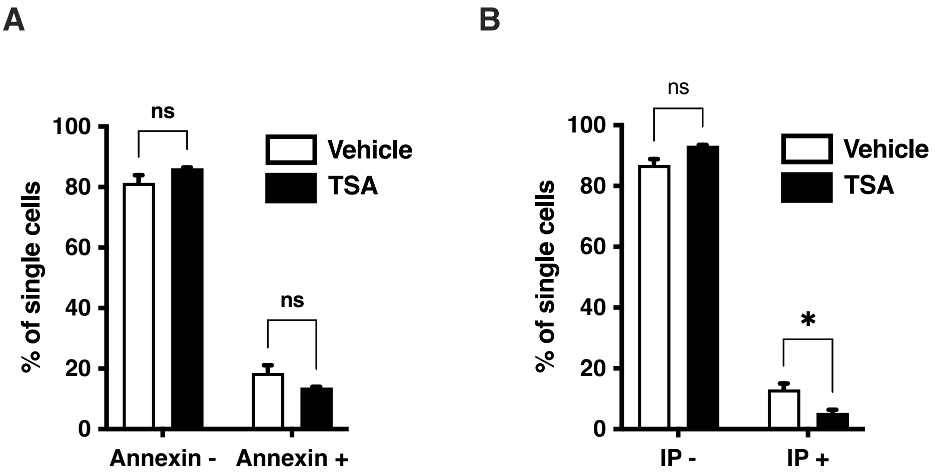

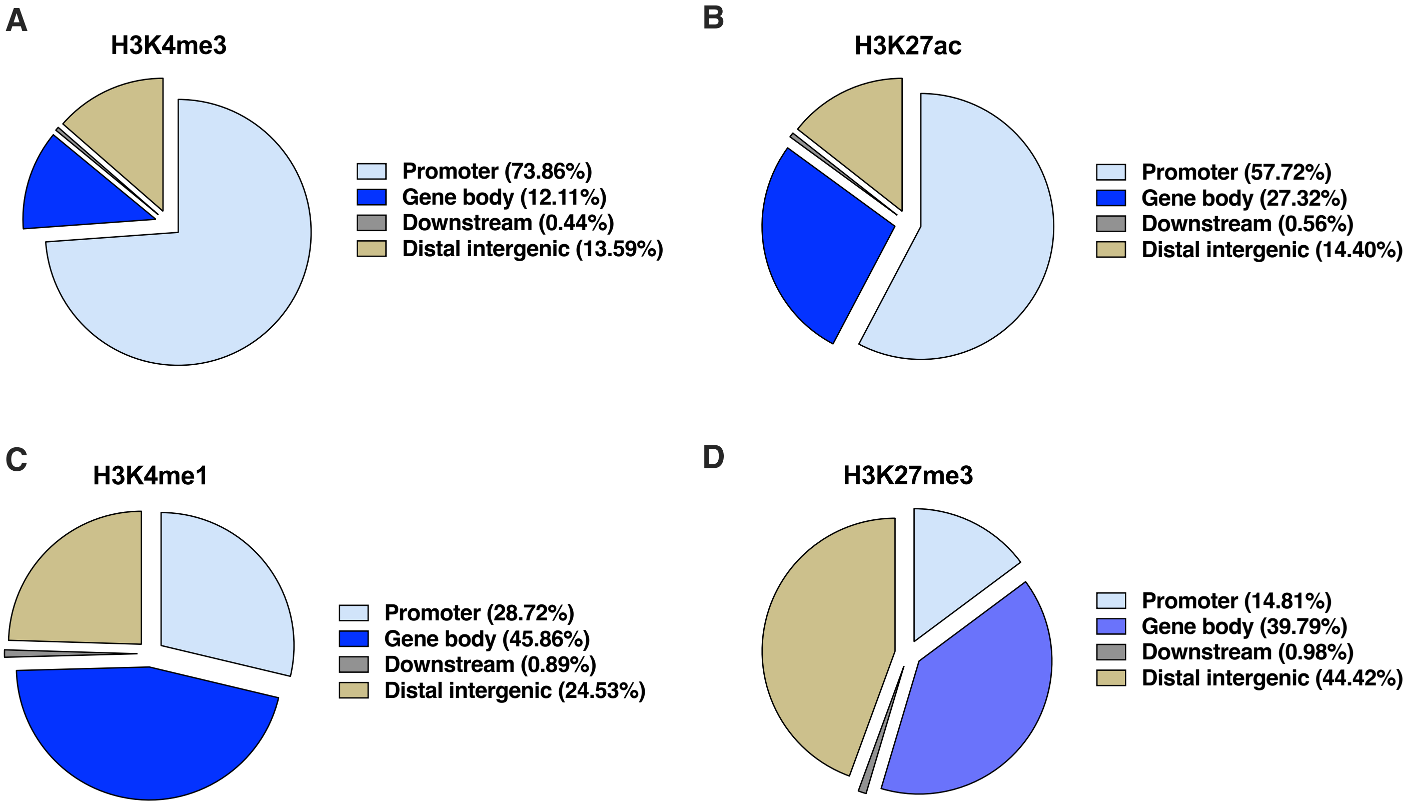

Oger *et al.*, Supplementary Figure 3

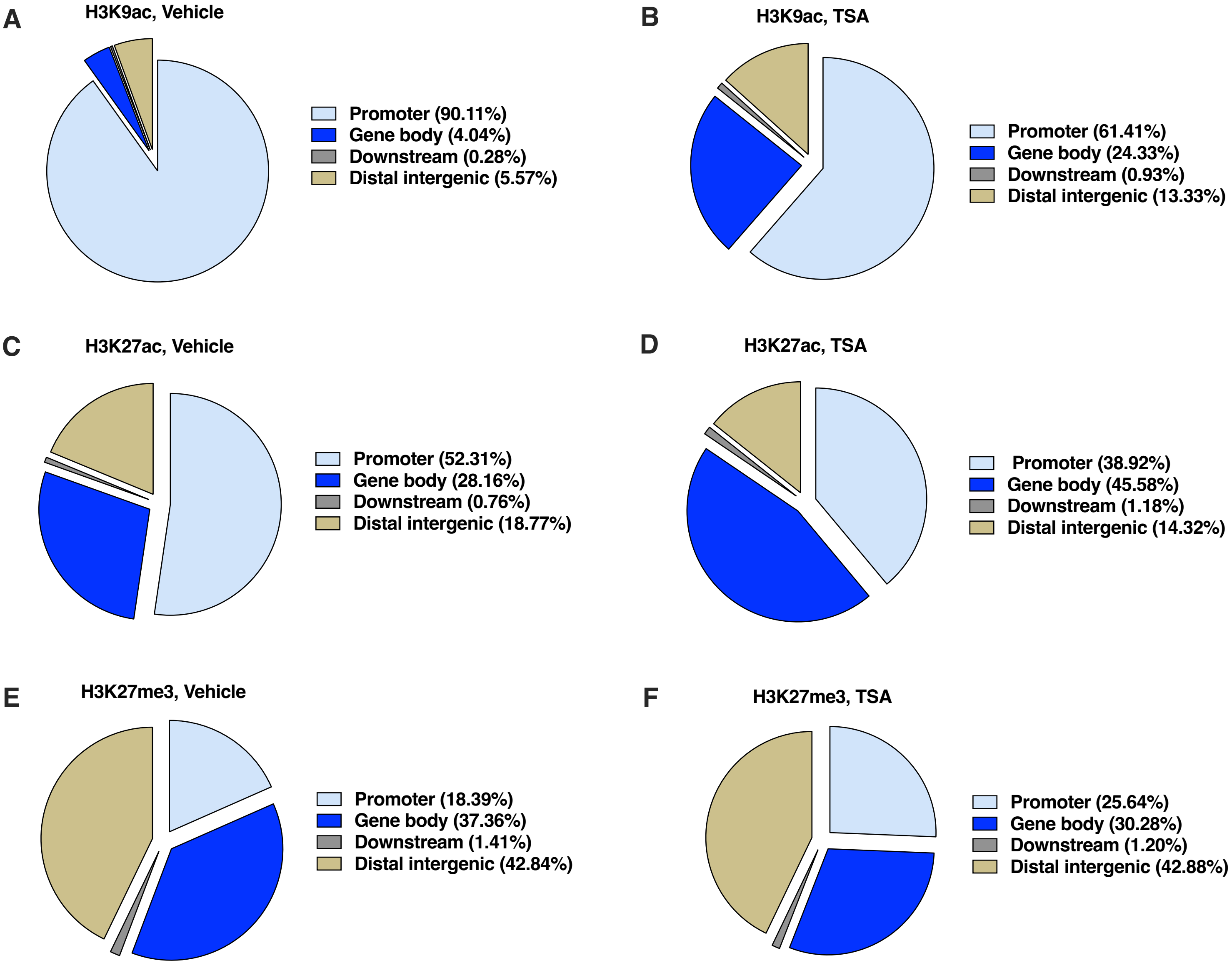

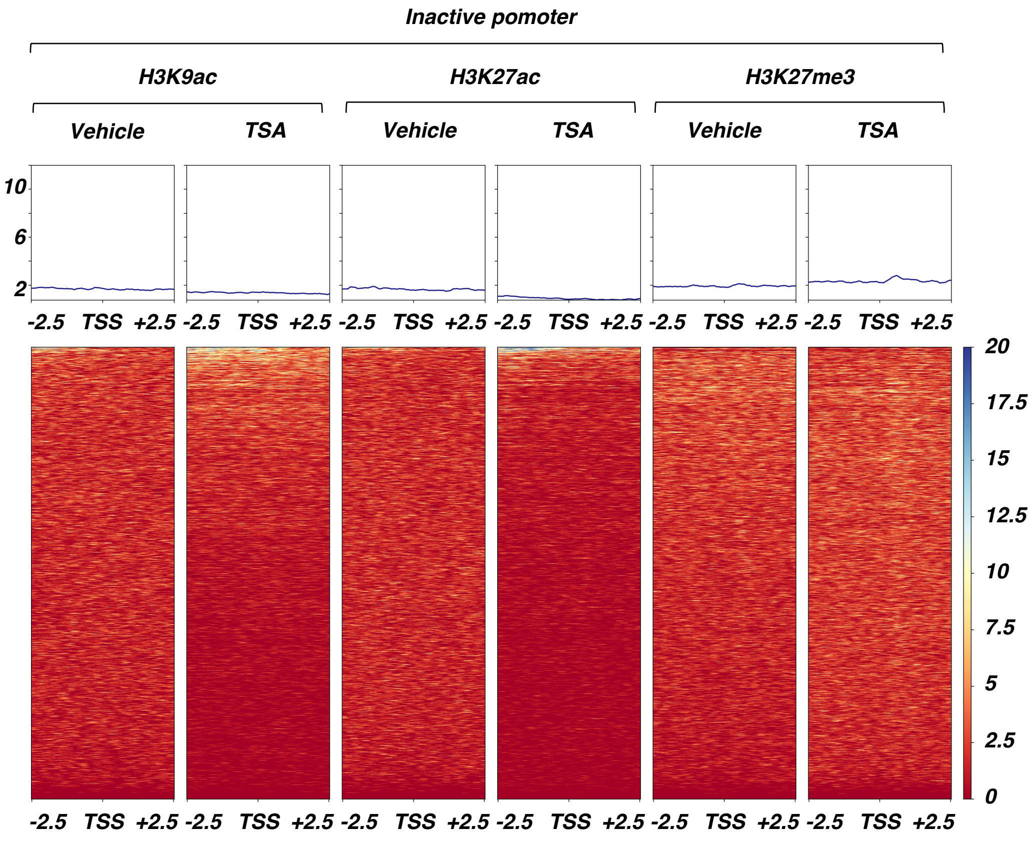

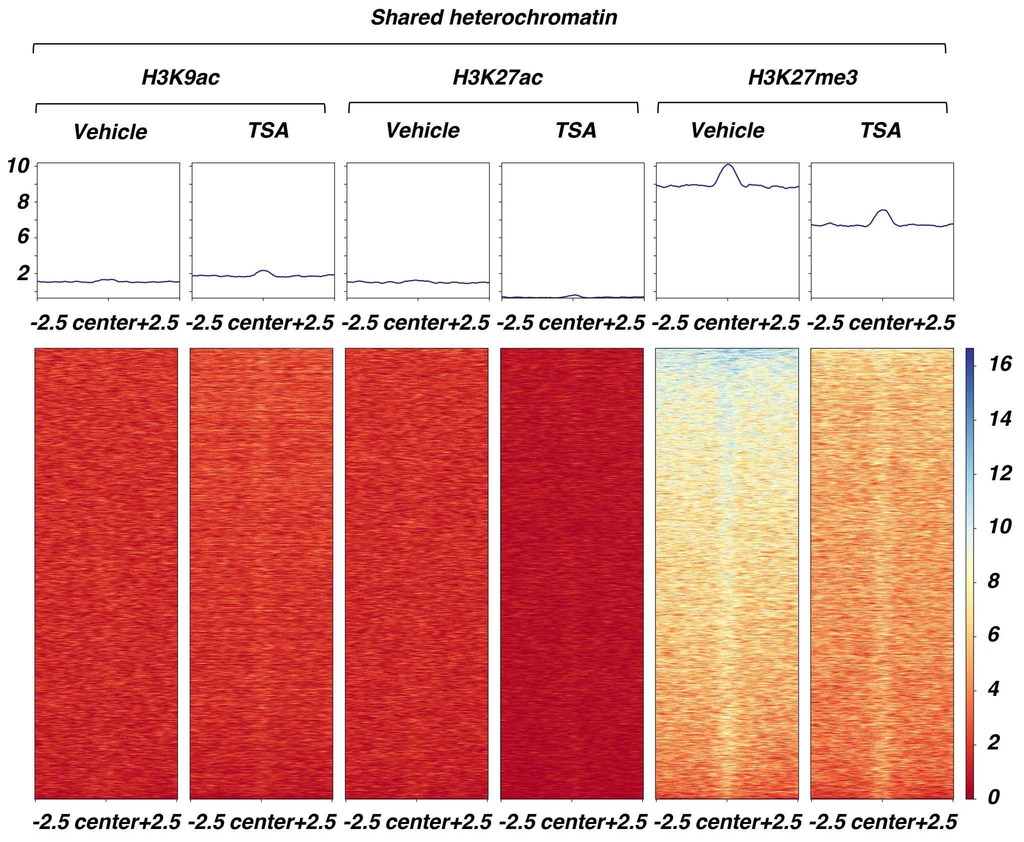

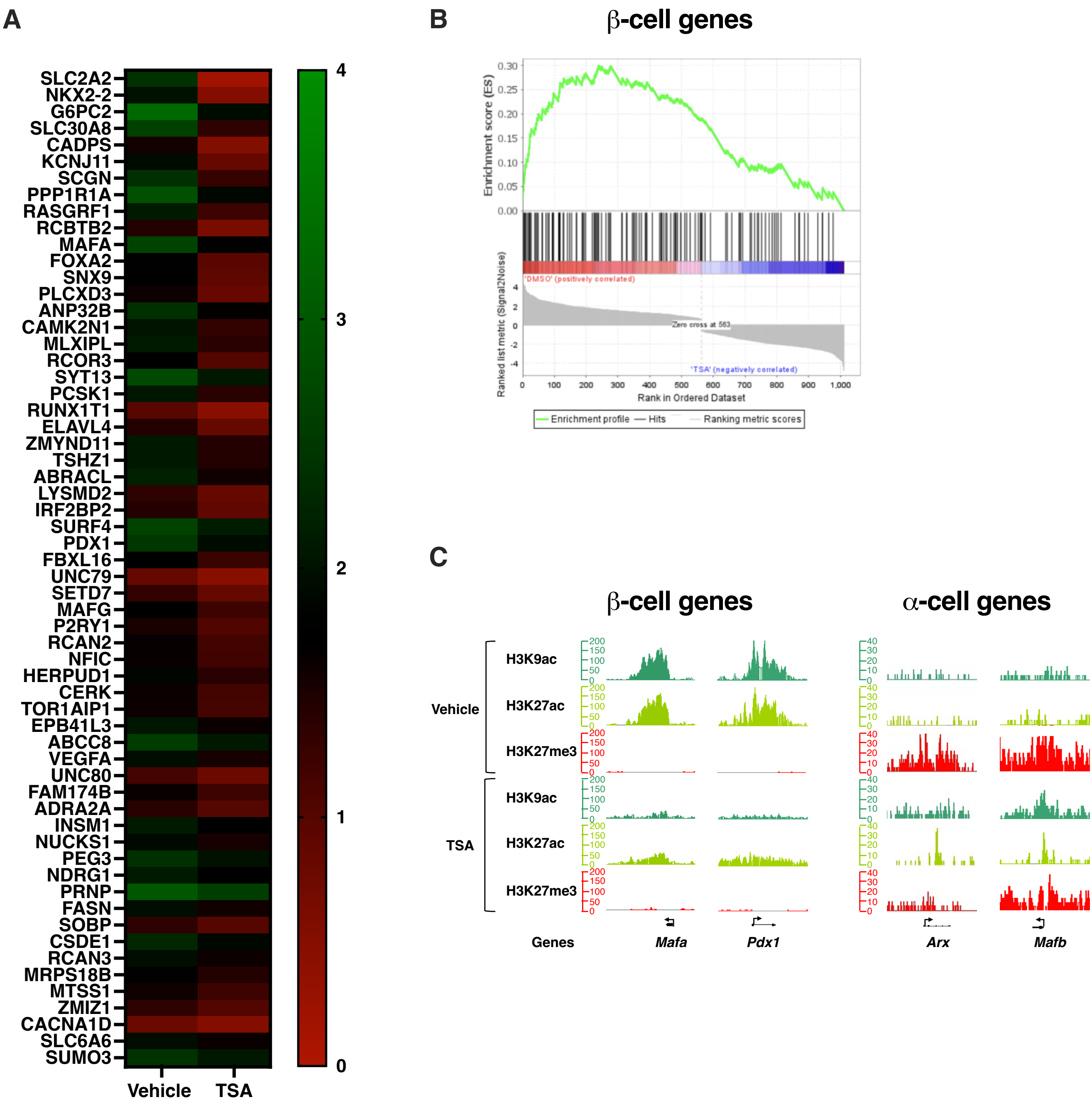
